# Phosphorylation-Coupled Autoregulation Maintains Functional ER Exit Sites

**DOI:** 10.1101/2025.06.18.660491

**Authors:** Miharu Maeda, Masashi Arakawa, Masaki Wakabayashi, Yukie Komatsu, Kota Saito

## Abstract

Secretory proteins are synthesized in the endoplasmic reticulum (ER) and begin their transport from specialized domains on the ER called ER exit sites (ERES). We previously demonstrated that the interaction between TANGO1 and Sec16 is critical for ERES formation. In this study, we reveal that the phosphorylation of TANGO1 and Sec16 is regulated by a FAM83A/CK1α-mediated negative feedback loop. Conversely, their dephosphorylation is regulated in a spatially distinct manner by different phosphatase complexes: PPP6R3/PPP6C for Sec16 and PPP1R15B/PPP1C for TANGO1. Excessive phosphorylation of either TANGO1 or Sec16 leads to ERES disassembly, while excessive dephosphorylation impairs secretion. Our findings demonstrate that maintaining a balanced phosphorylation state of TANGO1 and Sec16 through autoregulation by FAM83A/CK1α and the phosphatases PP1 and PP6 is essential for sustaining proper secretory activity at the ERES.

## INTRODUCTION

The secretory pathway is a fundamental cellular process in which proteins synthesized in the endoplasmic reticulum (ER) are transported through the Golgi apparatus and ultimately delivered to the extracellular space. Approximately one-third of all proteins synthesized in the cell are transported via this pathway. The budding of proteins from the ER is mediated by COPII factors at specialized ER subdomains known as ER exit sites (ERES). The COPII machinery is highly conserved from Saccharomyces cerevisiae to humans and has traditionally been considered essential for the formation of COPII-coated vesicles. The classical model describes a process in which the small GTPase Sar1 is activated by its guanine nucleotide exchange factor (GEF) Sec12, leading to its recruitment to the ER membrane. Subsequently, the inner coat proteins Sec23/Sec24 and outer coat proteins Sec13/Sec31 assemble to form COPII-coated vesicles, which then transport cargo proteins through the ER-Golgi intermediate compartment (ERGIC) to the Golgi apparatus^1^.

However, recent super-resolution microscopy studies have suggested that ERES in mammalian cells may not always generate discrete vesicles. Instead, a model has been proposed in which tubular structures extend from ERES toward the ERGIC, facilitating cargo transport^2, 3, 4^. Despite this structural variation, COPII factors remain essential for secretion from ERES, underscoring the importance of elucidating the precise roles of ERES-associated factors in mammalian cells.

Our previous studies demonstrated that the formation of ERES in mammalian cells requires the interaction between TANGO1 and Sec16^5^. Sec16 functions as a scaffold for COPII factors, recruiting cytoplasmic components to ERES^6^, while TANGO1, which is anchored to the ER membrane, serves as a scaffold for recruiting other ER-localized ERES factors, including cTAGE5 and Sec12^7, 8, 9^. The interaction between TANGO1 and Sec16 is, therefore, crucial for proper ERES assembly^10^. Furthermore, we previously reported that TANGO1 regulates the disassembly of ERES during mitosis^11^. Specifically, the phosphorylation state of TANGO1 is dynamically regulated throughout the cell cycle, and its phosphorylation during mitosis weakens its interaction with Sec16, leading to ERES disassembly. CK1δ and CK1ε act as the kinases responsible for TANGO1 phosphorylation, while PP1 functions as the phosphatase that dephosphorylates TANGO1. During mitosis, the reduced activity of PP1 results in a relative increase in TANGO1 phosphorylation, promoting ERES disassembly. However, the regulatory mechanisms controlling the activities of either kinase or phosphatase have remained unclear. In this study, we demonstrate that CK1α at the ER exit sites forms a complex with FAM83A, and that its kinase activity is spatially regulated, allowing phosphorylation of the TANGO1 and Sec16/Sec13 complex in interphase cells. We also show that PPP1R15B, a regulatory subunit of PP1 localized to the ER membrane, is responsible for dephosphorylating TANGO1, whereas PPP6R3, which is localized in the cytoplasm, functions as a regulatory subunit of PP6 and binds to Sec16 to promote its dephosphorylation. These findings reveal the presence of an autoregulatory mechanism that maintains ERES in a secretion-competent state through the spatially coordinated activities of kinases and phosphatases.

## RESULTS

### Sec13 is required for Sec16 dimerization and localization to ERES

Previous studies suggested that the interaction between Sec16 and Sec13 is required during the early stages of COPII vesicle formation^12^; however, the details of this process remain unclear. To investigate this further, we first examined the state of Sec16 upon Sec13 expression. As shown in Fig. 1A top panel, co-expression with Sec13 induced a band shift in Sec16, indicating a potential modification. Given that several post-translational modifications of Sec16 have been reported, we first examined whether phosphorylation was responsible for this shift. To test this, we performed Phos-tag blot analysis on immunoprecipitated Sec16 fractions. The results showed a marked increase in Sec16 phosphorylation upon Sec13 expression (Fig. 1A). Next, we examined the phosphorylation status of Sec16 in cells with Sec13 knockdown. In FLAG-Sec16 (1101aa-CT) stable cell lines, Sec13 depletion led to a significant reduction in Sec16 phosphorylation (Fig. 1B). A reduction in the apparent molecular weight of endogenous full-length Sec16 was also observed upon Sec13 knockdown, suggesting that phosphorylation of full-length Sec16 is similarly suppressed in the absence of Sec13 (see Fig. 1C, lysate Sec16). Sec16 has been reported to form homodimers^13^, and in control knockdown cells, endogenous full-length Sec16 was co-immunoprecipitated with FLAG-Sec16 (1101 aa-CT)(Fig. 1C). However, in Sec13-depleted cells, the amount of endogenous Sec16 co-precipitated with FLAG-Sec16 was markedly reduced. These findings strongly suggest that Sec13 is not only required for Sec16 phosphorylation but also plays a crucial role in Sec16 homodimerization. Next, we examined the localization of Sec16 and other ERES-associated factors in Sec13-depleted cells. The results showed that the ERES-localized Sec16 signal was markedly reduced upon Sec13 knockdown (Fig. S1). However, since the total protein levels of Sec16 remained largely unchanged (see Fig. 1C), we hypothesized that Sec16 might be dispersed throughout the cytoplasm. To test this, we performed PFA fixation, which is better suited for staining cytoplasmic components in GFP-Sec16 stable cell lines. As a result, Sec16 was found to be widely dispersed throughout the cytoplasm in Sec13-depleted cells (Fig. 1D). In contrast, the localization of TANGO1 remained largely unchanged, maintaining its association with ERES (Fig. 1D and S1). Notably, the small fraction of Sec16 that remained punctate exhibited strong colocalization with TANGO1, suggesting that it was still localized to ERES (Fig. 1D and S1). These findings strongly suggest that Sec13 is essential for Sec16 phosphorylation, dimerization, and localization to ERES.

**Fig. 1.**
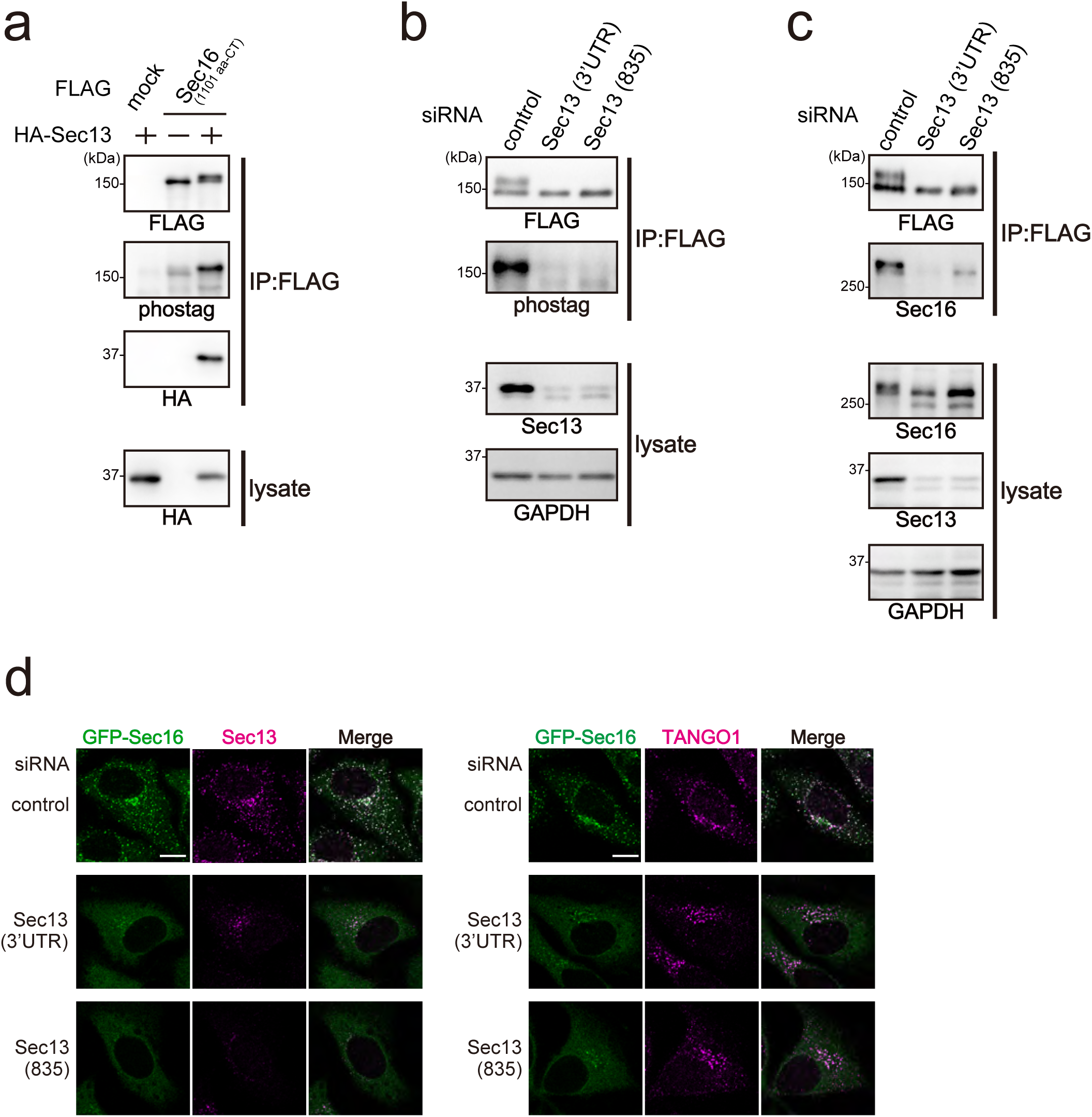
Sec13 is required for the phosphorylation, dimerization, and correct localization of Sec16. (A) 293T cells were transfected with indicated FLAG-tagged Sec16 (1101aa–CT) with HA-Sec13 constructs as indicated. Cell lysates were immunoprecipitated with anti-FLAG antibody and eluted with a FLAG peptide. Eluates were then subjected to SDS-PAGE followed by biotinylated Phos-tag detection or western blotting with anti-FLAG and anti-HA antibodies. Lysates were subjected to SDS-PAGE followed by western blotting with anti-HA antibody. (B) Doxycycline-inducible HeLa cells expressing FLAG-tagged Sec16 (1101aa–CT) were transfected with indicated siRNAs. Cell lysates were immunoprecipitated with anti-FLAG antibody and eluted with a FLAG peptide. Eluates were subjected to SDS-PAGE followed by biotinylated Phos-tag detection or western blotting with an anti-FLAG antibody. Lysates were subjected to SDS-PAGE followed by western blotting with anti-Sec13 and anti-GAPDH antibodies. (C) Doxycycline-inducible HeLa cells expressing FLAG-tagged Sec16 (1101aa–CT) were transfected with indicated siRNAs. Cell lysates were immunoprecipitated with anti-FLAG antibody and eluted with a FLAG peptide. Eluates were subjected to SDS-PAGE followed by western blotting with anti-FLAG and anti-Sec16-N antibodies. Lysates were subjected to SDS-PAGE followed by western blotting with anti-Sec16-N, anti-Sec13 and anti-GAPDH antibodies. (D) Doxycycline-inducible HeLa cells expressing GFP-tagged Sec16 were transfected with indicated siRNAs. Cells were then fixed with PFA and stained with anti-Sec13 or anti-TANGO1-CT antibodies. Bars, 10 µm.

### CK1α phosphorylates Sec16 in a Sec13-Dependent Manner

To identify kinases that phosphorylate Sec16 in a Sec13-dependent manner, we screened for factors interacting with Sec16. Specifically, we performed mass spectrometry analysis on proteins present in Sec16 immunoprecipitates from HeLa cells using an anti-Sec16 antibody, as well as FLAG immunoprecipitates from a stable FLAG-Sec16-expressing cell line. Additionally, we conducted a TurboID proximity labeling assay for Sec13, enriched biotinylated proteins using streptavidin beads, and identified them by mass spectrometry. We further supplemented our dataset with reported Sec16 interactors from the BIOGRID database and generated a Venn diagram integrating these four datasets (Fig. 2A). Among the 12 proteins identified across all datasets, we detected Sec23A, Sec23B, all four isoforms of Sec24, and Sec13, confirming the reliability of our screening approach. Notably, CSNK1A1(CK1α) was the only kinase present in this core set. In addition to CK1α, we selected 38 candidates based on their occurrence across multiple screening methods or their relevance to our kinase cDNA library. To assess their potential involvement in Sec16 phosphorylation, we co-expressed each candidate with Sec16 in 293T cells and analyzed Sec16 phosphorylation status using Phos-tag. As a result, CK1α, which received the highest score in Fig. 2A, was found to strongly enhance Sec16 phosphorylation (Fig. S2). Moreover, as shown in Fig. 2B, Sec16 phosphorylation by CK1α was further enhanced in the presence of Sec13. To further validate this, FLAG-Sec16 (1101 aa–CT) stably expressing cells were treated with CK1α siRNAs, and the immunoprecipitated proteins were subjected to an in vitro kinase assay. While Sec16 was efficiently phosphorylated in control knockdown cells, Sec16 immunoprecipitated from CK1α-depleted cells exhibited significantly reduced phosphorylation (Fig. 2C). Taken together, these results demonstrate that CK1α functions as a kinase that phosphorylates Sec16. Sec16 possesses an ERES localization domain (ELD), which is required for its interaction with TANGO1 and plays a crucial role in Sec16 localization to ERES ^5^. Sec16 also contains a central conserved domain (CCD) involved in Sec16-Sec16 interactions and a C-terminal conserved domain (CTCD) that mediates binding to Sec23^6^. Previous studies have reported that Sec13 interacts with Sec16 in the region between the ELD and CCD domains (Fig. 2D)^12^. To further investigate the interaction sites of CK1α and Sec13 on Sec16, we mapped the CK1α-binding region within Sec16. These results revealed that the CK1α-binding region of Sec16 is largely confined to the ELD, while the Sec13-binding region is located between the ELD and CCD, consistent with previous reports (Fig. 2E). Furthermore, the phosphorylation sites on Sec16 were found to be mainly concentrated within the ELD region (Fig. 2E). We previously identified CK1δ and CK1ε as kinases that phosphorylate TANGO1 during mitosis, leading to a weakened interaction between TANGO1 and Sec16^11^. To determine whether CK1δ and CK1ε also contribute to Sec13-dependent Sec16 phosphorylation, we next examined their effects. As shown in Fig. 2F, CK1α expression significantly increased Sec16 phosphorylation, whereas CK1δ and CK1ε had little effect. Furthermore, when CK1α was co-expressed with Sec13, Sec16 phosphorylation was further enhanced. In contrast, co-expression of CK1δ or CK1ε with Sec13 did not significantly increase Sec16 phosphorylation beyond the level observed with Sec13 alone (Fig. 2F). These findings indicate that CK1α, rather than CK1δ or CK1ε, functions as the kinase responsible for Sec16 phosphorylation in a Sec13-dependent manner.

**Fig. 2.**
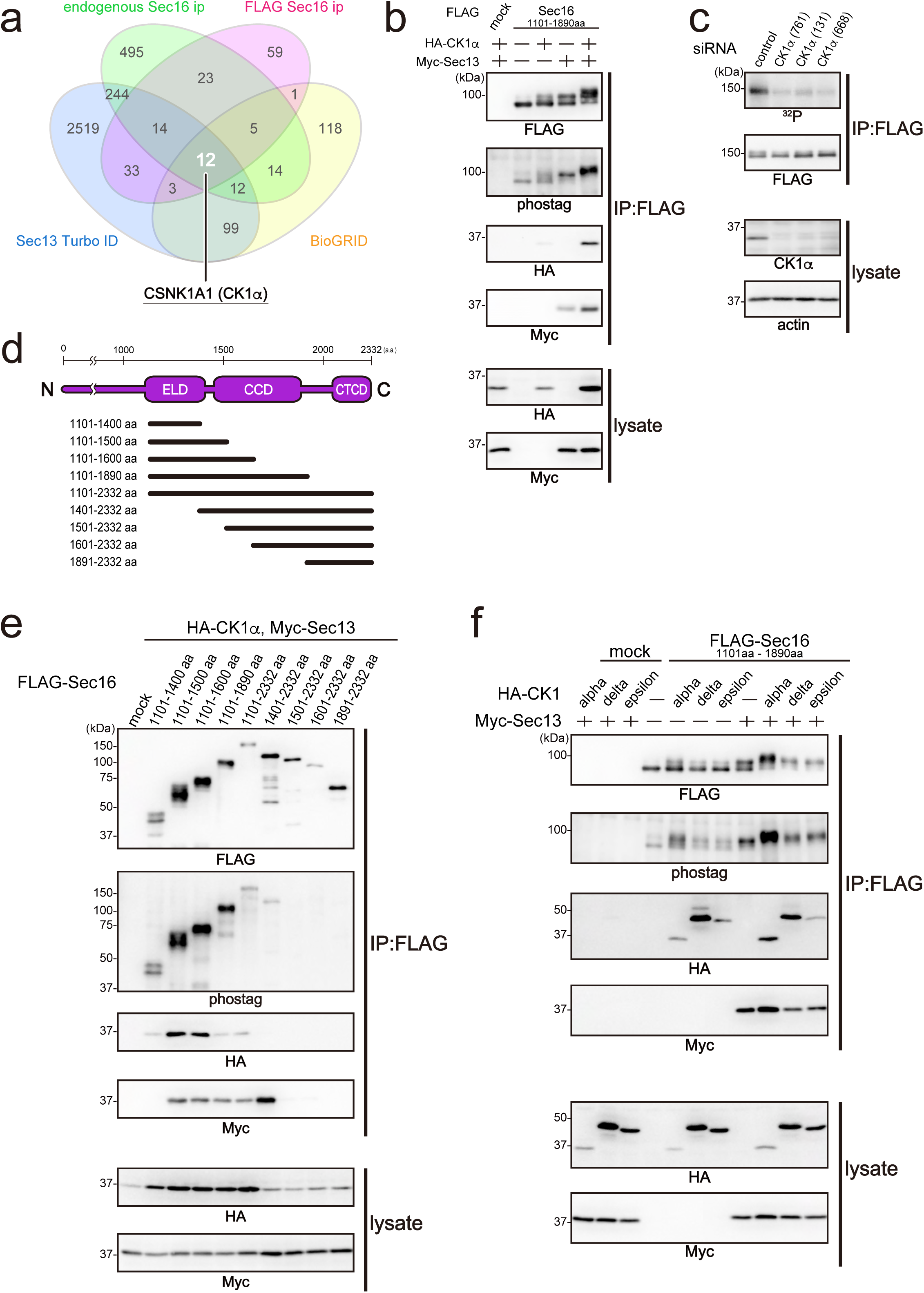
CK1α mediates Sec16 phosphorylation in a Sec13-dependent manner. (A) Venn diagram showing the overlap of Sec16-associated proteins identified by four approaches: immunoprecipitation of endogenous Sec16 from HeLa cells, FLAG-Sec16 from a stable cell line, TurboID-based proximity labeling with TurboID-Sec13, and curated interactors from the BIOGRID database. Numbers within the diagram indicate the number of proteins identified in each overlapping category. Twelve proteins were detected by all four methods, including core COPII components, validating the screening strategy. CSNK1A1 (CK1α) was the only kinase among them. (B) 293T cells were transfected with indicated FLAG-tagged Sec16 (1101–1890 aa) with HA-CK1α, Myc-Sec13 constructs as indicated. Cell lysates were immunoprecipitated with anti-FLAG antibody and eluted with a FLAG peptide. Eluates were then subjected to SDS-PAGE followed by biotinylated Phos-tag detection or western blotting with anti-FLAG, anti-HA, and anti-Myc antibodies. Lysates were subjected to SDS-PAGE followed by western blotting with anti-HA and anti-Myc antibodies. (C) Doxycycline-inducible HeLa cells expressing FLAG-tagged Sec16 (1101aa–CT) were transfected with indicated siRNAs. Cell lysates were immunoprecipitated with anti-FLAG antibody and eluted with a FLAG peptide. Eluates were then incubated with [γ-^32^P] ATP for in vitro kinase assay and then subjected to SDS-PAGE followed by exposing with phosphor plate analyzed by Typhoon FLA9500 or western blotting with anti-FLAG antibody. Lysates were subjected to SDS-PAGE followed by western blotting with anti-CK1α and anti-actin antibodies. (D) Schematic representation of human Sec16 domain organization. ELD, ERES localization domain; CCD, central conserved domain; CTCD, C-terminal conserved domain. (E) 293T cells were transfected with indicated FLAG-tagged Sec16 deletion constructs with HA-CK1α, Myc-Sec13 constructs as indicated. Cell lysates were immunoprecipitated with anti-FLAG antibody and eluted with a FLAG peptide. Eluates were then subjected to SDS-PAGE followed by biotinylated Phos-tag detection or western blotting with anti-FLAG, anti-HA, and anti-Myc antibodies. Lysates were subjected to SDS-PAGE followed by western blotting with anti-HA and anti-Myc antibodies. (F) 293T cells were transfected with HA-CK1α or HA-CK1δ or HA-CK1ε with FLAG-tagged Sec16 (1101–1890 aa) and Myc-Sec13 constructs as indicated. Cell lysates were immunoprecipitated with anti-FLAG antibody and eluted with a FLAG peptide. Eluates were then subjected to SDS-PAGE followed by biotinylated Phos-tag detection or western blotting with anti-FLAG, anti-HA, and anti-Myc antibodies. Lysates were subjected to SDS-PAGE followed by western blotting with anti-HA and anti-Myc antibodies.

### CK1-mediated Sec16 phosphorylation promotes its dynamics and efficient secretion from ERES

To identify the phosphorylation sites on Sec16 modified by CK1α in the presence of Sec13, we performed phosphorylation proteomics analysis on Sec16 (1101-1890aa) phosphorylated under these conditions (Fig. 3A). This analysis revealed that 14 serine and threonine residues were potential phosphorylation sites. To further validate their functional significance, we generated non-phosphorylatable SA mutants, in which the identified phosphorylation sites were substituted with alanine, and performed an in vitro kinase assay. FLAG-Sec16 (1101 aa-CT) stable cell lines were also generated for the SA mutant, and FLAG immunoprecipitates from these cells were subjected to an in vitro kinase assay. As shown in Fig. 3B, while phosphorylation was detected in cells expressing wild-type Sec16, it was markedly suppressed in the SA mutant. These results allowed us to identify specific phosphorylation sites on Sec16 that are modified in a Sec13-dependent manner. To investigate the impact of Sec16 phosphorylation on its localization, we generated GFP-Sec16 stable cell lines expressing the wild-type (WT), SA mutant, and phosphomimetic SE mutant and analyzed their localization. The results showed that all Sec16 variants colocalized with TANGO1, indicating that Sec16 localizes to ERES regardless of its phosphorylation status (Fig. S3). Next, to determine whether Sec16 phosphorylation affects its turnover rate at ERES, we performed fluorescence recovery after photobleaching (FRAP) analysis to assess their membrane dynamics. The results showed that both the wild-type and SE mutant exhibited similar recovery rates after photobleaching, whereas the SA mutant displayed a significantly delayed recovery (Fig. 3C and 3D). These findings suggest that Sec16 phosphorylation enhances its turnover at the ERES membrane. The comparable FRAP recovery rates between the wild-type and SE mutant further imply that Sec16 undergoes efficient phosphorylation under steady-state conditions. Next, we examined the impact of Sec16 phosphorylation on secretion from ERES. To assess this, we evaluated the transport of ManII using a RUSH assay in stable cell lines expressing either wild-type or SA mutant 3×FLAG-Sec16. As shown in Fig. 3E, in cells expressing wild-type Sec16, a majority of ManII relocated to the Golgi apparatus within 20 minutes after biotin addition. In contrast, in cells expressing the SA mutant, ManII was occasionally observed in the reticular ER. The quantification shown in Fig. 3F further confirmed that ManII transport was significantly delayed in SA-expressing cells compared to wild-type cells.

**Fig. 3.**
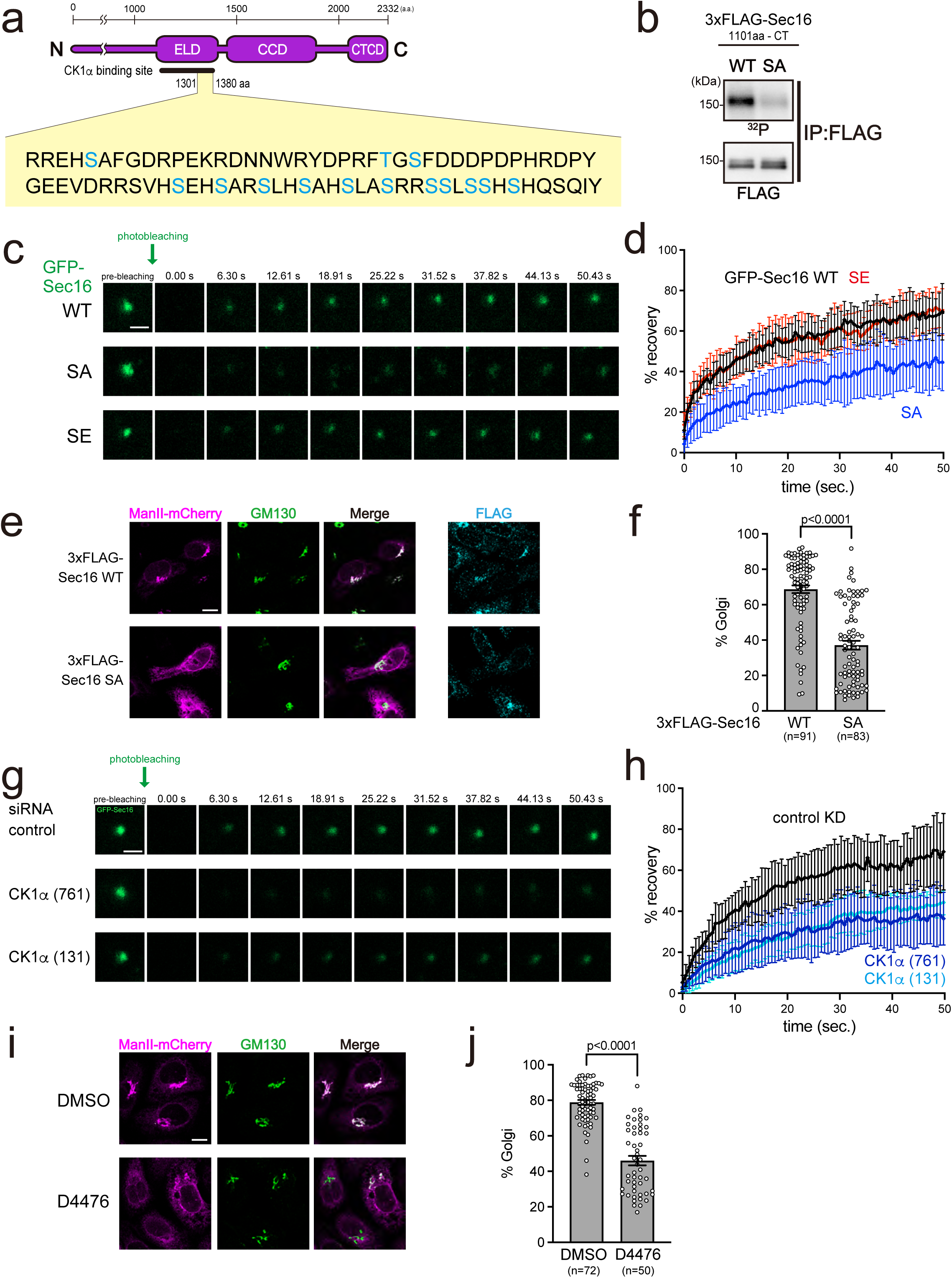
CK1α-mediated phosphorylation of Sec16 regulates its turnover and secretion from ERES. (A) Phosphorylation sites on Sec16 identified by phosphoproteomic analysis in the presence of Sec13. A total of 14 serine and threonine residues were detected as candidate phosphorylation sites. (B) Doxycycline-inducible HeLa cells expressing either FLAG-Sec16 (1101aa–CT) or FLAG-Sec16 (1101aa–CT) SA were extracted and cell lysates were immunoprecipitated with anti-FLAG antibody and eluted with a FLAG peptide. Eluates were then incubated with [γ-^32^P] ATP for in vitro kinase assay and then subjected to SDS-PAGE followed by exposing with phosphor plate analyzed by Typhoon FLA9500 or western blotting with anti-FLAG antibody. (C) Doxycycline-inducible HeLa cells expressing either GFP-Sec16 or GFP-Sec16 SA or GFP-Sec16 SE were subjected to FRAP assay. Representative images are shown before bleaching (pre-bleaching) and at the indicated time points after photobleaching. Bars, 1 µm. (D) Quantification of fluorescence recovery from FRAP assays as in (C). FRAP curves show the mean fluorescence intensity of 10 individual regions and error bars indicate the S.D. Fluorescence values were normalized to the pre-bleach intensity. (E) Doxycycline-inducible HeLa cells expressing either 3xFLAG-Sec16 or 3xFLAG-Sec16 SA mutant were transfected with Str-KDEL_ManII-SBP-mCherry. RUSH chase was initiated with the addition of biotin and terminated after 20 minutes with cold methanol fixation. The cells were then stained with anti-GM130 and anti-FLAG antibodies. Bars, 10 μm. (F) Quantification of the percentage of ManII-mCherry signals in the GM130-positive region out of all ManII-mCherry signals for (E). Data were analyzed by two-tailed Student’s t test. (G) Doxycycline-inducible HeLa cells expressing GFP-Sec16 were transfected with indicated siRNAs and subjected to FRAP assay. Representative images are shown before bleaching (pre-bleaching) and at the indicated time points after photobleaching. Bars, 1 µm. (H) Quantification of fluorescence recovery from FRAP assays as in (G). FRAP curves show the mean fluorescence intensity of 10 individual regions and error bars indicate the S.D. Fluorescence values were normalized to the pre-bleach intensity. (I) Doxycycline-inducible HeLa cells expressing Str-KDEL/ManII-SBP-mCherry were treated with or without 100 µM D4476 for 4 h. RUSH chase was started with the addition of biotin and fixed after 20 min with cold methanol. The cells were stained with an anti-GM130 antibody. Bars, 10 μm. (J) Quantification of the percentage of ManII-mCherry signals in the GM130-positive region out of all ManII-mCherry signals for (I). Data were analyzed by two-tailed Student’s t test.

Next, to determine whether Sec16 phosphorylation affecting its turnover rate and secretion is directly mediated by CK1α in cells, we measured the turnover rate of Sec16 using FRAP in CK1α-depleted cells. The results showed that compared to control knockdown, CK1α knockdown significantly reduced the turnover rate of Sec16 (Fig. 3G and 3H). A similar reduction was also observed when cells were treated with D4476, a CK1α inhibitor, suggesting that CK1α-mediated phosphorylation of Sec16 is essential for its steady-state turnover at ERES (Fig. S4A and S4B). Furthermore, we performed a RUSH assay in D4476-treated cells and found that ManII transport was delayed upon CK1α inhibition (Fig. 3I and 3J). Taken together, these results strongly suggest that CK1α-mediated phosphorylation of Sec16 regulates its turnover rate at ERES under steady-state conditions and plays a crucial role in secretion from ERES.

### FAM83A acts as a scaffold for CK1α-mediated phosphorylation of Sec16

CK1α is known to regulate various cellular functions, including membrane trafficking, cell cycle progression, apoptosis, autophagy, and circadian rhythm, and has numerous identified substrates. Although CK1α is constitutively active in biochemical assays, the mechanisms governing its activity regulation remain largely unknown^14^. Recent studies have reported that FAM83 family proteins interact with CK1 through their DUF1669 domain, contributing to its localization to various intracellular compartments^15, 16^. Based on this, we sought to identify a FAM83 protein involved in Sec16 phosphorylation by examining the intracellular localization of CK1α in the presence of different FAM83 family members. We co-expressed FAM83A, FAM83D, FAM83G, and FAM83H with CK1α in cells and analyzed their localization (Fig. 4A). Consistent with previous reports, CK1α exhibited strong colocalization with all tested FAM83 proteins, localizing to various intracellular compartments (Fig. 4A). When compared with Sec16 localization, however, only FAM83A-expressing cells exhibited partial colocalization of FAM83A and CK1α with Sec16 (Fig. 4A). Given that kinase-substrate interactions are often transient and weaken upon phosphorylation, we examined the colocalization of Sec16 with a kinase-dead CK1α mutant (CK1α K46A) in the presence of FAM83A. As shown in Fig. 4B, CK1α K46A and FAM83A strongly colocalized with Sec16. In contrast, FAM83D, FAM83G and FAM83H failed to colocalize with Sec16 when co-expressed with CK1α K46A (Fig. S5). These results suggest that FAM83A may be involved in Sec16 phosphorylation. To further investigate the interactions among Sec16, CK1α, and FAM83A, we performed Sec16 immunoprecipitation in cells expressing CK1α K46A and FAM83A, either individually or together. The results revealed that FAM83A binds to Sec16, and the amount of CK1α K46A associated with Sec16 increased in the presence of FAM83A (Fig. 4C, lanes 3 and 4). Next, we mapped the FAM83A binding region on Sec16 and found that, similar to CK1α, FAM83A binds to the ELD domain of Sec16 (Fig. 4D). These findings suggest that CK1α interacts with Sec16 through FAM83A. To further examine the effect of CK1α on the FAM83A-Sec16 interaction, we performed FAM83A immunoprecipitation in cells co-expressing FAM83A, Sec16 WT or SA mutant, and either wild-type or kinase-dead CK1α. The results showed that the interaction between wild-type Sec16 and FAM83A was significantly reduced in the presence of wild-type CK1α, but not in the presence of kinase-dead CK1α (Fig. 4E). Furthermore, when using the Sec16 SA mutant, which cannot be phosphorylated by CK1α, the addition of wild-type CK1α did not significantly reduce the interaction between FAM83A and Sec16 (Fig. 4E). These findings indicate that phosphorylation of Sec16 by CK1α weakens its interaction with FAM83A. Therefore, FAM83A likely functions as a scaffold that facilitates CK1α-mediated phosphorylation of Sec16.

**Fig. 4.**
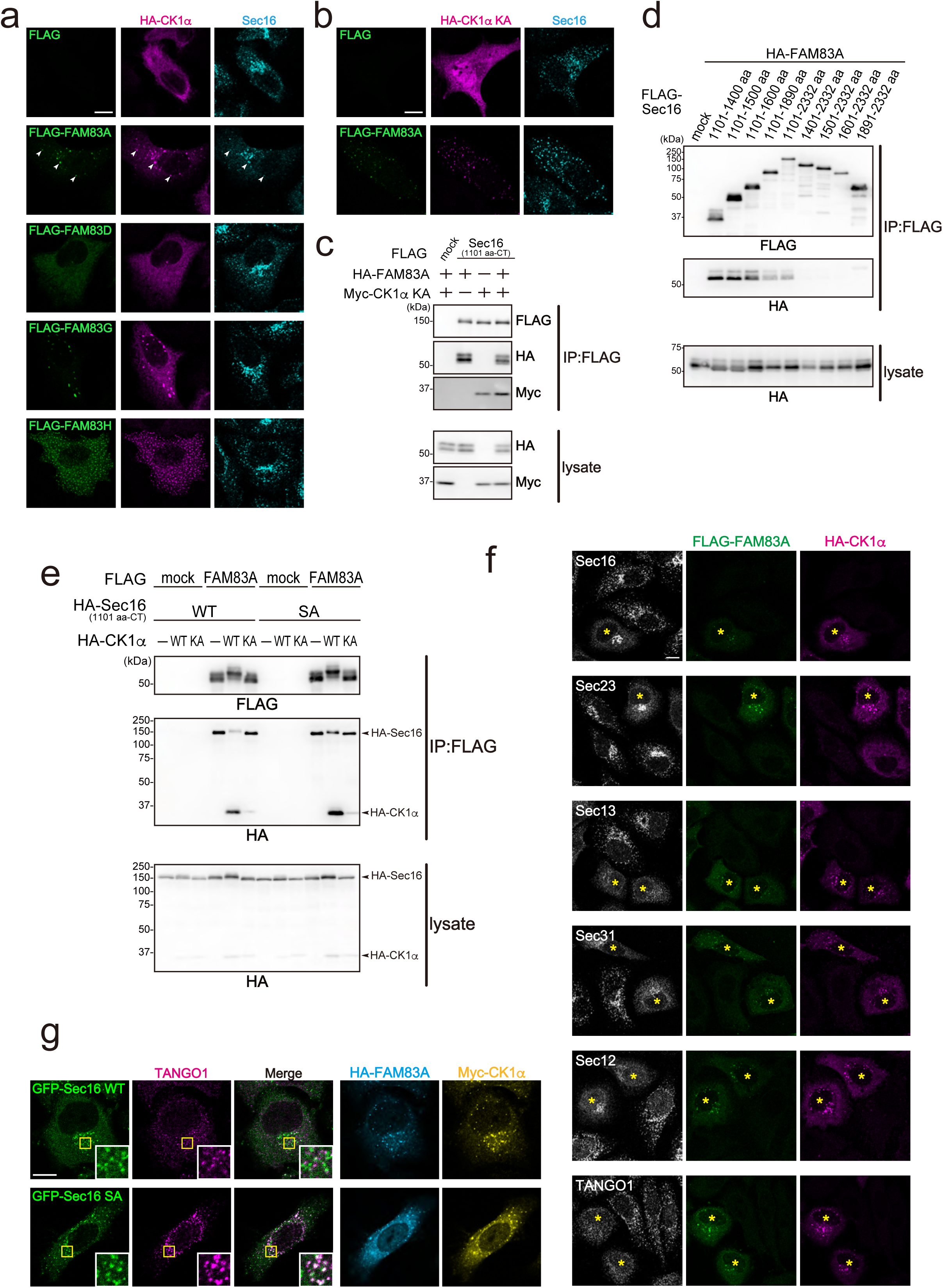
FAM83A serves as a scaffold for CK1α-driven phosphorylation of Sec16. (A) HeLa cells transfected with FLAG-tagged FAM83 family proteins as indicated with HA-CK1α were fixed and stained with anti-Sec16-C, anti-FLAG, and anti-HA antibodies. Bars, 10 µm. (B) HeLa cells transfected with FLAG-FAM83A with HA-CK1α K46A were fixed and stained with anti-Sec16-C, anti-FLAG, and anti-HA antibodies. Bars, 10 µm. (C) 293T cells were transfected with FLAG-tagged Sec16 (1101aa–CT) with HA-FAM83A and Myc-CK1α K46A constructs as indicated. Cell lysates were immunoprecipitated with anti-FLAG antibody and eluted with a FLAG peptide. Eluates were then subjected to SDS-PAGE followed by western blotting with anti-FLAG, anti-HA, and anti-Myc antibodies. Lysates were subjected to SDS-PAGE followed by western blotting with anti-HA and anti-Myc antibodies. (D) 293T cells were transfected with indicated FLAG-tagged Sec16 deletion constructs with HA-FAM83A constructs as indicated. Cell lysates were immunoprecipitated with anti-FLAG antibody and eluted with a FLAG peptide. Eluates were then subjected to SDS-PAGE followed by western blotting with anti-FLAG and anti-HA antibodies. Lysates were subjected to SDS-PAGE followed by western blotting with anti-HA antibody. (E) 293T cells were transfected with FLAG-FAM83A and HA-Sec16 (1101aa–CT) or HA-Sec16 (1101aa–CT) SA and HA-CK1α or HA-CK1α K46A as indicated. Cell lysates were immunoprecipitated with anti-FLAG antibody and eluted with a FLAG peptide. Eluates were then subjected to SDS-PAGE followed by western blotting with anti-FLAG and anti-HA antibodies. Lysates were subjected to SDS-PAGE followed by western blotting with anti-HA antibody. (F) HeLa cells transfected with FLAG-FAM83A with HA-CK1α were fixed and stained with anti-FLAG, anti-HA, anti-Sec16-C (rabbit) or anti-Sec23 or anti-Sec13 or anti-Sec31 or anti-Sec12 or anti-TANGO1-CT antibodies. Bars, 10 µm. (G) Doxycycline-inducible HeLa cells expressing either GFP-Sec16 or GFP-Sec16 SA were transfected with HA-FAM83A and Myc-CK1α. Cells were fixed and stained with anti-HA, anti-Myc, anti-TANGO1-CT antibodies. Bars, 10 µm.

### Excessive phosphorylation of Sec16 by CK1α leads to the disassembly of ERES

To investigate the effects of FAM83A and CK1α on ERES organization, we analyzed the localization of various ERES components in cells co-expressing both proteins. The results showed that the expression of FAM83A and CK1α caused the dispersion of cytoplasmic ERES factors, and even membrane-associated ERES proteins such as Sec12 and TANGO1 were distributed throughout the ER (Fig. 4F). To determine whether this dispersion of ERES components is caused by CK1α-mediated phosphorylation of Sec16, we examined the effect of FAM83A and CK1α co-expression in cells expressing either wild-type Sec16 or the Sec16-SA mutant. As shown in Fig. 4G, co-expression of FAM83A and CK1α in cells expressing wild-type Sec16 resulted in the dispersal of both Sec16 and TANGO1, consistent with previous observations. However, in cells expressing the Sec16-SA mutant, not only Sec16 and TANGO1 but also FAM83A and CK1α remained co-localized at ERES, and ERES disassembly was suppressed (Fig. 4G). These findings suggest that while moderate phosphorylation of Sec16 by CK1α is required to promote ERES turnover and support secretion, excessive phosphorylation disrupts ERES integrity. Thus, a balanced level of CK1α-mediated phosphorylation of Sec16 appears to be critical for maintaining functional ERES (further discussed in the Discussion section).

### Proper phosphorylation of TANGO1 is critical for ERES function and efficient secretion

We previously reported that during mitosis, TANGO1 is phosphorylated by CK1δ and CK1ε, isoforms of CK1, and that this phosphorylation disrupts the interaction between TANGO1 and Sec16, leading to ERES disassembly^11^. Based on this, we hypothesized that the FAM83A–CK1α complex may also be involved in the phosphorylation of TANGO1, in addition to Sec16. To test this, we examined the effect of co-expression of FAM83A and CK1α on TANGO1 phosphorylation. Interestingly, the expression of FAM83A and CK1α alone did not significantly alter the phosphorylation state of TANGO1 (Fig. 5A, lanes 2 and 3). However, the addition of Sec16 led to a marked increase in TANGO1 phosphorylation (Fig. 5A, lanes 2 and 5). Furthermore, the interaction between TANGO1 and Sec16 was significantly reduced upon co-expression of FAM83A and CK1α. These results indicate that FAM83A–CK1α promotes TANGO1 phosphorylation in a Sec16-dependent manner.

**Fig. 5.**
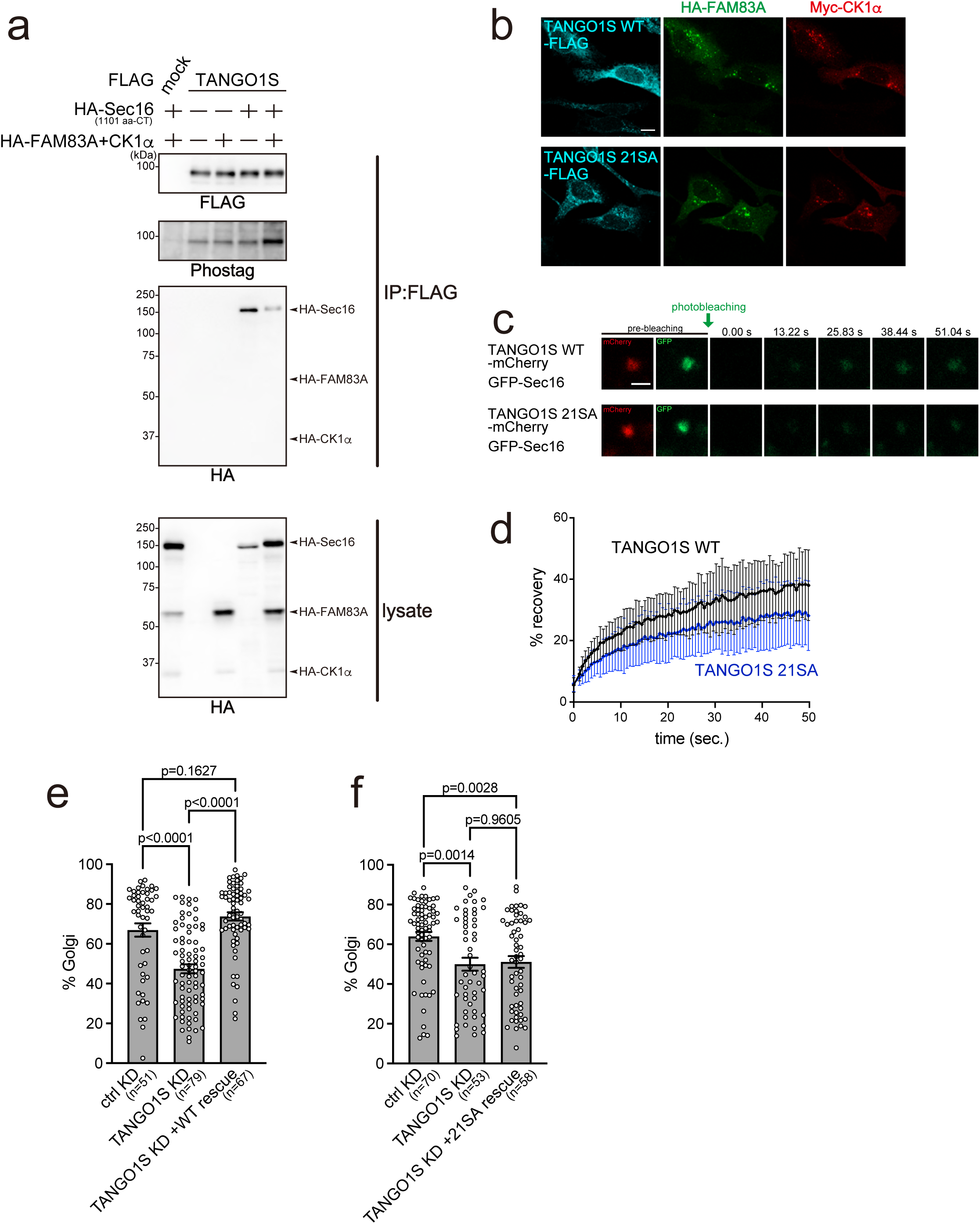
Proper phosphorylation of TANGO1 by the FAM83A–CK1α complex is essential for ER exit site function. (A) 293T cells were transfected with TANGO1S-FLAG with HA-Sec16 (1101 aa-CT), HA-FAM83A and HA-CK1α constructs as indicated. Cell lysates were immunoprecipitated with anti-FLAG antibody and eluted with a FLAG peptide. Eluates were then subjected to SDS-PAGE followed by biotinylated Phos-tag detection or western blotting with anti-FLAG and anti-HA antibodies. Lysates were subjected to SDS-PAGE followed by western blotting with anti-HA antibody. (B) Doxycycline-inducible HeLa cells expressing either TANGO1S-FLAG or TANGO1S 21SA-FLAG were transfected with HA-FAM83A and Myc-CK1α. Cells were fixed and stained with anti-FLAG, anti-HA, anti-Myc antibodies. Bars, 10 µm. (C) Doxycycline-inducible HeLa cells expressing GFP-Sec16 were transfected with either TANGO1S WT-mCherry or TANGO1S 21SA-mCherry and subjected to FRAP assay. Representative images are shown before bleaching (pre-bleaching) and at the indicated time points after photobleaching. Bars, 1 µm. (D) Quantification of fluorescence recovery from FRAP assays as in (C). FRAP curves show the mean fluorescence intensity of 10 individual regions and error bars indicate the S.D. Fluorescence values were normalized to the pre-bleach intensity. (E and F) Doxycycline-inducible HeLa cells expressing either TANGO1S-FLAG (E) or TANGO1S SA mutant-FLAG (F) were transfected with the indicated siRNAs. The cells were treated with or without doxycycline and transfected with Str-KDEL_ManII-SBP-mCherry. RUSH chase was initiated with the addition of biotin and terminated after 20 min with cold methanol fixation. The cells were stained with anti-GM130 and anti-FLAG antibodies. Quantification of percentage of ManII-mCherry signals in the GM130-positive region out of all ManII-mCherry signals. Data were analyzed by two-way ANOVA and Tukey’s multiple comparison test. Representative images are shown in Supplemental Fig. 6.

Next, we examined whether the disruption of ERES induced by FAM83A and CK1α expression could be suppressed by the expression of a non-phosphorylatable TANGO1 mutant^11^. In cells stably expressing wild-type TANGO1, FAM83A and CK1α caused TANGO1 to disperse throughout the ER. In contrast, this dispersal was suppressed in cells expressing the phosphorylation-deficient TANGO1 21SA mutant (Fig. 5B). To further investigate the state of ERES in these cells, we performed a FRAP assay using GFP-Sec16. The results showed that Sec16 turnover was significantly slower in TANGO1 21SA-expressing cells compared to those expressing wild-type TANGO1 (Fig. 5C and D). We then performed a RUSH assay using inducible cell lines stably expressing either wild-type TANGO1S or the TANGO1S 21SA mutant. Knockdown of endogenous TANGO1S in the absence of induction of the rescue constructs resulted in delayed secretion in both cell lines (Fig. S6A and S6B, Fig. 5E and F). When expression of either wild-type or 21SA TANGO1S was induced and endogenous TANGO1S was selectively silenced, secretion was restored in cells rescued with wild-type TANGO1S but remained impaired in cells rescued with the 21SA mutant (Fig. S6A and S6B, Fig. 5E and F). Taken together, these results demonstrate that excessive phosphorylation of TANGO1 weakens its interaction with Sec16 and leads to ERES disassembly, whereas insufficient phosphorylation also impairs secretion. Therefore, proper regulation of TANGO1 phosphorylation is essential for maintaining functional ERES and efficient protein secretion.

### The ER-localized PP1 regulatory subunit PPP1R15B is involved in the dephosphorylation of TANGO1

While this study has provided new insights into the phosphorylation control of TANGO1 and Sec16, the regulatory mechanism of TANGO1 dephosphorylation by PP1 remains unclear. Given that nearly 200 PP1 regulatory subunits have been reported, we performed TurboID proximity labeling using TANGO1S to identify candidate PP1 regulatory subunits located in proximity to TANGO1S^17^. Among the candidates, PPP1R15B showed the highest score in the TurboID dataset and is known to localize to the ER^18^, making it a strong candidate for further analysis. Co-immunoprecipitation assays in HEK293T cells co-expressing TANGO1S with either PPP1R12A or PPP1R15B revealed that only PPP1R15B specifically bound to TANGO1S (Fig. 6A). To further validate the role of PPP1R15B, we knocked down its expression in a TANGO1S-stable cell line and analyzed the phosphorylation status of TANGO1S. Knockdown of PPP1R15B led to a clear increase in TANGO1S phosphorylation (Fig. 6B). In parallel, we generated a phospho-specific antibody that recognizes phosphorylated Sec16. This antibody successfully detected the reduction of Sec16 phosphorylation following Sec13 knockdown (Fig. S7). Using this antibody, we examined Sec16 phosphorylation following PPP1R15B knockdown. In contrast to TANGO1S, the phosphorylation of Sec16 remained unchanged (Fig. 6C), indicating substrate specificity. Taken together with previous observations that the TANGO1 21SE mutant is diffusely localized across the ER^11^, these results suggest a model in which phosphorylated TANGO1S diffuses throughout the ER, where it is recognized and dephosphorylated by PPP1R15B/PPP1C complex.

**Fig. 6.**
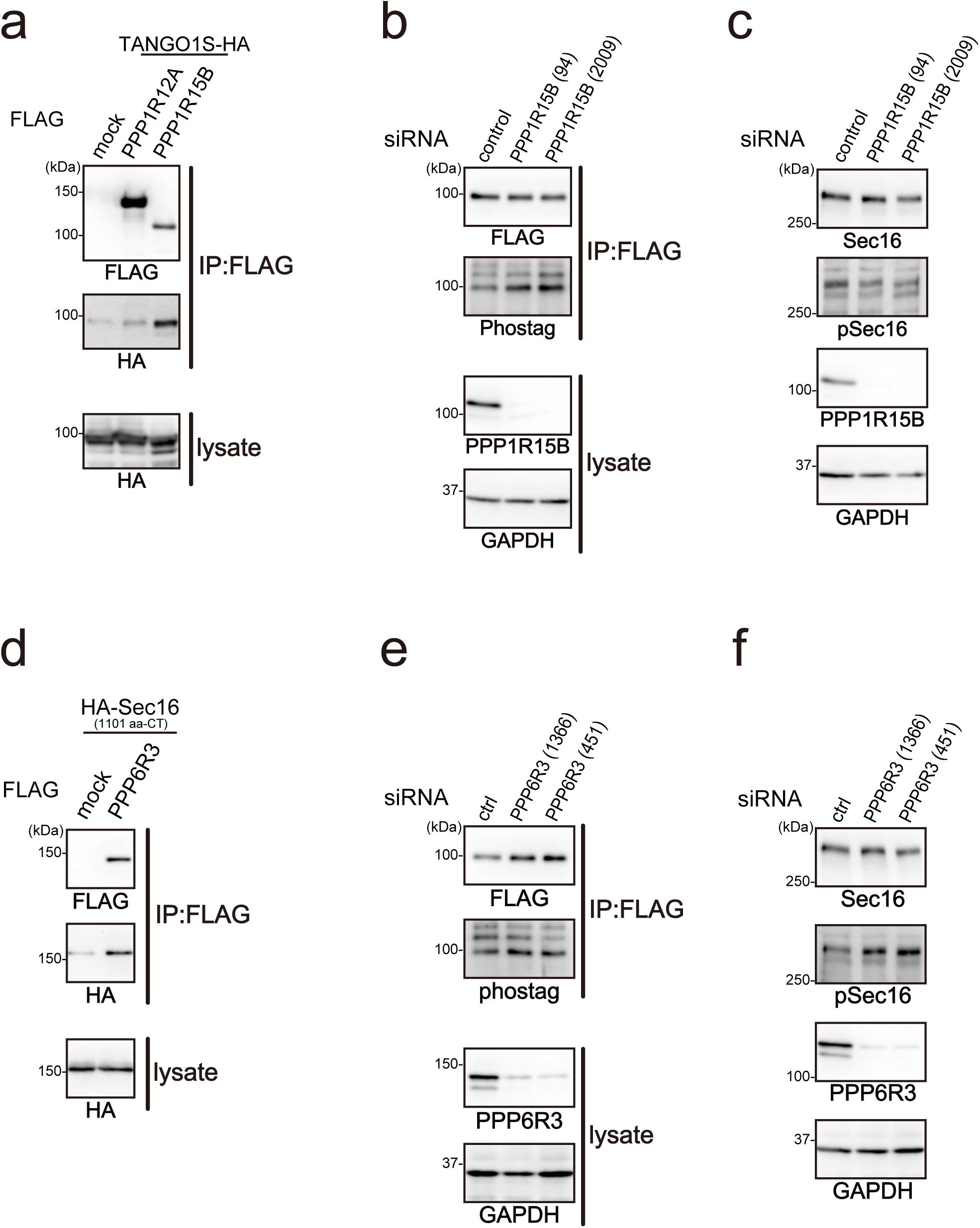
PP1 and PP6 regulate TANGO1 and Sec16 dephosphorylation through their respective regulatory subunits. (A) 293T cells were transfected with FLAG-tagged PPP1R indicated constructs with TANGO1S-HA. Cell lysates were immunoprecipitated with anti-FLAG antibody and eluted with a FLAG peptide. Eluates were then subjected to SDS-PAGE followed by western blotting with anti-FLAG and anti-HA antibodies. Lysates were subjected to SDS-PAGE followed by western blotting with anti-HA antibody. (B) Doxycycline-inducible HeLa cells expressing TANGO1S-FLAG were transfected with indicated siRNAs. Cell lysates were immunoprecipitated with anti-FLAG antibody and eluted with a FLAG peptide. Eluates were then subjected to SDS-PAGE followed by biotinylated Phos-tag detection or western blotting with anti-FLAG antibody. Lysates were subjected to SDS-PAGE followed by western blotting with anti-PPP1R15B and anti-GAPDH antibodies. (C) HeLa cells were transfected with indicated siRNAs. Cell lysates were subjected to SDS-PAGE followed by western blotting with anti-Sec16-N, anti-pSec16, anti-PPP1R15B and anti-GAPDH antibodies. (D) 293T cells were transfected with FLAG-tagged PPP6R3 with HA-Sec16 (1101 aa-CT). Cell lysates were immunoprecipitated with anti-FLAG antibody and eluted with a FLAG peptide. Eluates were then subjected to SDS-PAGE followed by western blotting with anti-FLAG and anti-HA antibodies. Lysates were subjected to SDS-PAGE followed by western blotting with anti-HA antibody. (E) Doxycycline-inducible HeLa cells expressing TANGO1S-FLAG were transfected with indicated siRNAs. Cell lysates were immunoprecipitated with anti-FLAG antibody and eluted with a FLAG peptide. Eluates were then subjected to SDS-PAGE followed by biotinylated Phos-tag detection or western blotting with anti-FLAG antibody. Lysates were subjected to SDS-PAGE followed by western blotting with anti-PPP6R3 and anti-GAPDH antibodies. (F) HeLa cells were transfected with indicated siRNAs. Cell lysates were subjected to SDS-PAGE followed by western blotting with anti-Sec16-N, anti-pSec16, anti-PPP6R3 and anti-GAPDH antibodies.

### Cytoplasm-localized PPP6R3/PPP6C contributes to the dephosphorylation of Sec16

Since PPP1R15B/PPP1C did not contribute to Sec16 dephosphorylation, we next investigated whether PP6 is involved in Sec16 dephosphorylation, based on the mass spectrometry results identifying Sec16-associated proteins. Co-immunoprecipitation analysis confirmed an interaction between PPP6R3, a regulatory subunit of PP6, and Sec16 (Fig. 6D). We then assessed the phosphorylation status of TANGO1 and Sec16 following knockdown of PPP6R3. While Phos-tag analysis showed that the phosphorylation level of TANGO1 remained largely unchanged (Fig. 6E), the phosphorylation of Sec16 was markedly increased (Fig. 6F).

## DISCUSSION

We previously demonstrated that the interaction between TANGO1 and Sec16 is crucial for the formation of ERES^5^. In the present study, we identified CK1α as a kinase responsible for phosphorylating Sec16 and showed that this phosphorylation is regulated by FAM83A. Since the interaction between FAM83A and Sec16 is weakened upon Sec16 phosphorylation, this interaction likely functions as a negative feedback loop to prevent excessive Sec16 phosphorylation. Furthermore, we found that the phosphorylation of TANGO1 by FAM83A/CK1α requires the presence of Sec16, suggesting that once TANGO1 becomes hyperphosphorylated and dissociates from Sec16, this phosphorylation process is terminated. Thus, the phosphorylation of both TANGO1 and Sec16 by FAM83A/CK1α is finely tuned to avoid excessive modification (Fig. 7).

**Fig. 7.**
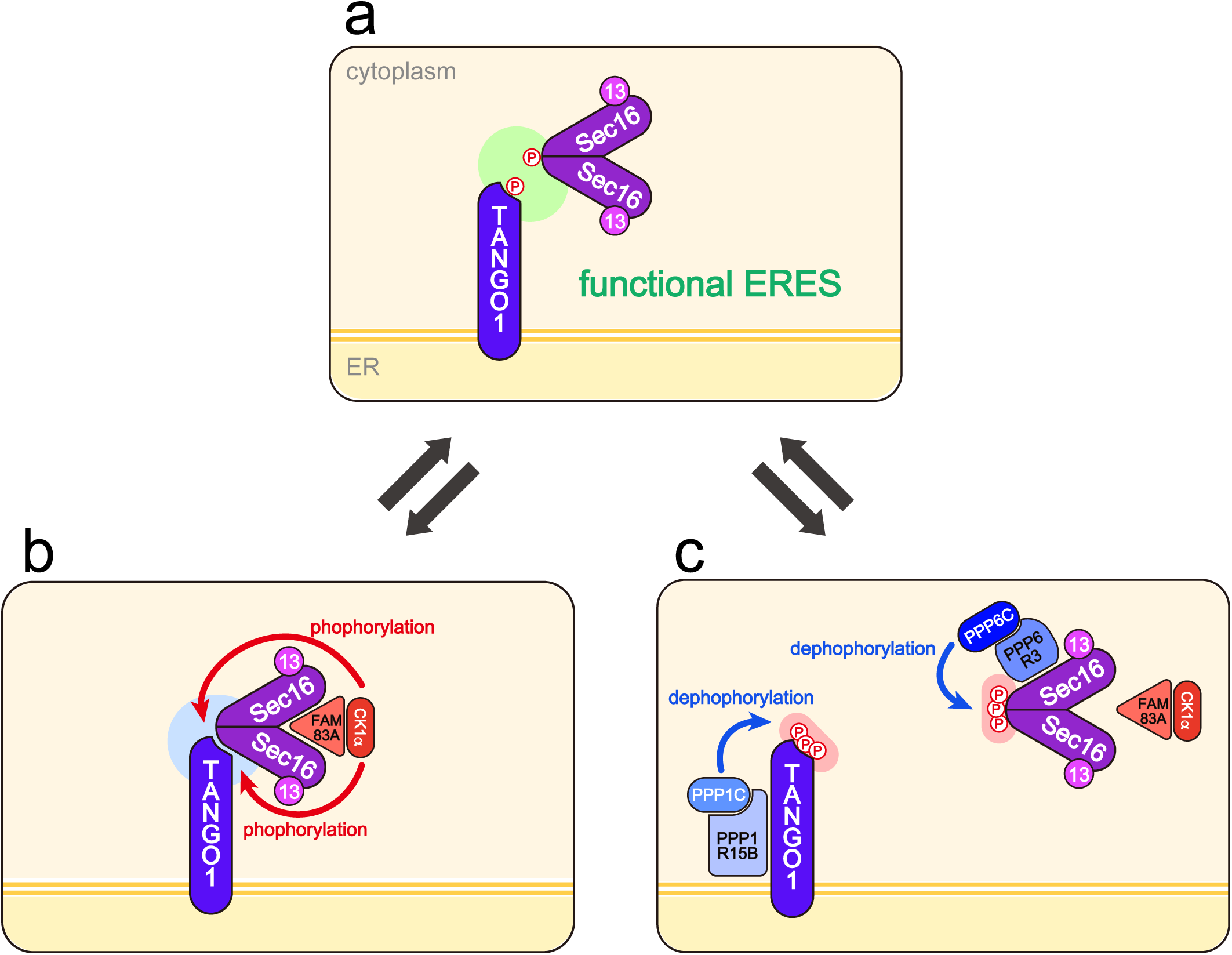
Model of ER exit site autoregulation. (A) A functional ERES is maintained by keeping TANGO1 and Sec16 in a moderately phosphorylated state. (B) To form a functional ERES, the FAM83A/CK1α complex binds to Sec16 and phosphorylates it. Once Sec16 is phosphorylated, FAM83A dissociates from Sec16. TANGO1 is also phosphorylated by FAM83A/CK1α, but this phosphorylation requires Sec16 to be bound to TANGO1. (C) Phosphorylated TANGO1 is dephosphorylated by the ER-localized PPP1R15B/PPP1C complex and returns to the ERES. Sec16 is dephosphorylated by the cytosolic PPP6R3/PPP6C complex.

Previously, we reported that hyperphosphorylated TANGO1 diffuses throughout the ER membrane^11^. In this study, we demonstrate that the phosphatase PPP1R15B/PPP1C, localized to the ER reticular membrane, can dephosphorylate TANGO1, enabling it to return to the ERES. Similarly, when Sec16 becomes hyperphosphorylated, it is dephosphorylated by PPP6R3/PPP6C, which are cytoplasmically localized. The dephosphorylated Sec16 can then re-bind to FAM83A and re-enter the phosphorylation cycle mediated by CK1α (Fig. 7). Together, these observations support a model in which ERES in interphase cells are maintained in a secretion-competent state via an autoregulatory mechanism involving a dynamic balance between phosphorylation by FAM83A/CK1α and dephosphorylation by PP1 and PP6, without requiring external stimuli (Fig. 7).

Importantly, we found that either excessive Sec16 phosphorylation promotes ERES disassembly or excessive Sec16 dephosphorylation disrupts secretion. Similarly, while TANGO1 hyperphosphorylation promotes its diffusion from ERES, its excessive dephosphorylation suppresses Sec16 turnover and delays secretion. Therefore, maintaining the phosphorylation balance of TANGO1 and Sec16 via autoregulation is critical for preserving ERES functionality (Fig. 7).

In this study, we initially isolated CK1α as a kinase activity dependent on Sec13. This finding suggests that the formation of a complex between Sec16 and Sec13 is important for proper secretion. In contrast to this stable complex, the interaction between TANGO1 and Sec16 appears to be more transient. In wild-type ER exit sites where TANGO1 and Sec16 interact, FRAP analysis revealed that the turnover rate of Sec16 is comparable to that of the phospho-mutant Sec16, indicating that Sec16 is rapidly exchanged. Therefore, rather than forming a stable complex, TANGO1 and Sec16 likely maintain their interaction in a dynamic, continuously exchanging manner.

DYRK3, previously identified as a kinase that phosphorylates Sec16, is thought to regulate secretion via the phase separation of Sec16^19^. Among the 14 phosphorylation sites we identified as CK1α targets on Sec16, five residues—within the range we could verify—overlap with sites phosphorylated by DYRK3. Additionally, Sec16 is known to exist in multiple transcript variants, and recent findings suggest that alternative splicing of Sec16 may influence COPII transport efficiency^20^. In our study, we used the full-length transcript variant X12 of Sec16 (6996 bp). In contrast, the DYRK3-related study employed an isoform initially cloned as “Sec16L,” as full-length, which corresponds to various C-terminal fragments found in multiple transcript variants (covering the sequence from amino acid 179 onward in transcript variant X12)^13^. Whether our findings and the phase separation regulated by DYRK3 are influenced by the specific transcript variant of Sec16 remains an intriguing question for future investigation. Although our results highlight FAM83A as a regulator of CK1α-mediated phosphorylation of Sec16, this regulation may still be compatible with a phase separation-based mechanism. Whether phase separation contributes to the newly proposed autoregulatory model or whether DYRK3, a kinase specialized for phase separation, is controlled by specific upstream stimuli remains an important question. It is also possible that DYRK3 activity is integrated into the FAM83A–CK1α, PP1, and PP6 autoregulatory axis, warranting further investigation.

We identified PP6 as a phosphatase responsible for dephosphorylating Sec16. PP6 was previously implicated in the dephosphorylation of COPII components in budding yeast and is known to interact with CK1δ in mammalian cells to regulate ER-to-Golgi transport^21^. Our identification of PP6 as a Sec16 phosphatase in mammalian cells suggests that PP6 may also dephosphorylate other COPII components localized at ERES. Interestingly, PPP6R3/PPP6C does not dephosphorylate TANGO1, while PPP1R15B/PPP1C dephosphorylates TANGO1 but not Sec16, suggesting that even among ERES-localized proteins, distinct phosphatases may be utilized depending on whether the substrate is membrane-anchored or cytoplasmic. In the aforementioned DYRK3 study, PP1 was suggested as a candidate for Sec16 phosphatase^19^. However, the inhibitors used in that study, such as okadaic acid and calyculin A, also inhibit PP6^22^. Moreover, the observed reduction in GFP-Sec16A turnover upon PPP1CC overexpression may reflect PPP1CC targeting TANGO1 phosphorylation, similar to what we observed in Fig. 5D. It will be important to clarify whether the phosphatases involved in Sec16 dephosphorylation differ depending on the upstream kinase, whether PP1 and PP6 function cooperatively, or whether only PP6 is involved. To date, three regulatory subunits of PP6 have been identified. Determining whether these are used selectively and how ANKRD family proteins, known to associate with PP6 complexes, contribute to specificity will be an important direction for future research^23^.

Previously, we reported that TANGO1 is phosphorylated by CK1δ and CK1ε during mitosis^11^. Here, we investigated whether CK1α also contributes to TANGO1 phosphorylation during mitosis. In cells with CK1δ and CK1ε knockdown, the co-localization of Sec16 and Sec31 during mitosis was significantly higher than in control knockdown cells, indicating partial inhibition of ERES disassembly. A similar phenotype was observed in CK1α knockdown cells (Fig. S8), indicating that CK1α also contributes to ERES disassembly during mitosis. Investigating how phosphorylation of TANGO1 by CK1 family kinases is regulated during mitosis, including the role of the FAM83 family, is a valuable future direction for research.

Sec16 has previously been shown to act as a sensor that transmits extracellular environmental signals, such as growth cues and nutrient deprivation, to the secretory machinery via phosphorylation and other modifications^24, 25, 26^. However, the regulation of Sec16 and TANGO1 phosphorylation revealed in this study is distinct from such stimulus-driven regulation. Instead, it is an autoregulatory mechanism that operates constitutively under steady-state conditions. The question of why cells maintain a system that continuously consumes energy to preserve an intermediate phosphorylation state of TANGO1 and Sec16 is intriguing. One possibility is that this system enables rapid responses to environmental changes or cargo load variations by keeping components primed for either increased phosphorylation or enhanced dephosphorylation. In this study, we elucidated for the first time an autoregulatory mechanism that maintains optimal phosphorylation of TANGO1 and Sec16, thereby creating a secretion-competent ERES during interphase. Future studies should address how this secretory environment adapts to various external signals, stresses, and changes in cargo levels.

## METHODS

### Antibodies

A female 6-wk-old Wistar rat (CLEA Japan, Inc.) was immunized with GST-tagged Sec12 (93–239 aa) in TiterMax Gold (TiterMax USA, Inc.). Splenocytes were fused with PAI mouse myeloma cells using Polyethylene Glycol 1500 (Roche). Hybridoma supernatants were screened by indirect ELISA with ColdTF-tagged Sec12 (93–239 aa) as the antigen. Positive hybridoma lines were subcloned, grown in serum-free medium (Nihon Pharmaceutical) supplemented with HT (Life Technologies), and purified with protein G–Sepharose (GE Healthcare)^8^. A female 6-wk-old Wistar rat (CLEA Japan) was immunized with FLAG-tagged Sec23A in TiterMax Gold (TiterMax USA). Splenocytes were fused with PAI mouse myeloma cells using polyethylene glycol (Roche). Hybridoma supernatants were screened by indirect ELISA with His-tagged Sec23A as the antigens. Positive hybridoma lines were subcloned, grown in serum-free medium (Nihon Pharmaceutical) supplemented with hypoxanthine-thymidine (Thermo Fisher Scientific), and purified with protein G-Sepharose (GE Healthcare)^5^. A female 8-wk-old Wistar rat (CLEA Japan) was immunized with KLH-conjugated Sec16-C (2,319–2,332 aa) in TiterMax Gold (TiterMax USA). Splenocytes were fused with PAI mouse myeloma cells using polyethylene glycol (Roche). Hybridoma supernatants were screened by indirect ELISA with Sec16-C peptide as the antigens. Positive hybridoma lines were subcloned, grown in serum-free medium (Nihon Pharmaceutical) supplemented with hypoxanthine-thymidine (Thermo Fisher Scientific), and purified with protein G-Sepharose (GE Healthcare). Polyclonal antibodies against Sec16-N (374–387 aa), Sec16-C (2,319–2,332 aa), and TANGO1-CT (1,884–1,898 aa for TANGO1L; 762–776 aa for TANGO1S) were raised in rabbits by immunization with keyhole limpet hemocyanin-conjugated peptides and affinity-purified by columns conjugated with the peptides (Thermo Fisher Scientific)^7, 9, 27, 28, 29^. Polyclonal antibodies against Sec13 were raised in rabbits by immunization with His_6_-Sec13 and affinity-purified by columns conjugated with GST-Sec13^30^. A phospho-specific antibody against Sec16 was generated by immunizing rabbits with a phosphorylated peptide corresponding to amino acids 1345–1358 of human Sec16, in which Ser1353 and Ser1356 are phosphorylated. The phosphopeptide was conjugated to a carrier protein and used for immunization. The resulting antiserum was affinity-purified using the phosphorylated peptide, followed by absorption with the corresponding non-phosphorylated peptide to remove antibodies recognizing the unphosphorylated form. Other antibodies were as follows: FLAG (mouse; SIGMA M2 for IP, WB, mouse; proteintech 66008-4-Ig for IF, rabbit; Cell signaling D6W5B, rat; agilent), HA (rat;Roche 3F10, rabbit; proteintech), Myc (mouse; 9E10), CK1α (rabbit; proteintech), actin (mouse; SIGMA), GAPDH (mouse; Santacruz), PPP1R15B (rabbit; proteintech) and Sec31 (mouse; BD).

### Cell culture and transfection

HeLa and 293T cells were cultured in DMEM supplemented with 10% fetal bovine serum. Lipofectamine RNAi max (Thermo Fisher Scientific) was used for transfecting siRNA. For plasmids transfection, polyethylenimine “MAX” (polysciences) or Fugene 4K (Promega) were used. Doxycycline-inducible stable HeLa cell lines expressing FLAG-Sec16, FLAG-Sec16 SA mutant, FLAG-Sec16 (1101 aa-CT), FLAG-Sec16 (1101aa–CT) SA mutant, GFP-Sec16, GFP-Sec16 SE mutant, GFP-Sec16 SA mutant, TANGO1S-FLAG, TANGO1S 21SA mutant-FLAG, Str-KDEL/ManII-SBP-mCherry were made described previously^5^. Proteins were induced by incubation with doxycycline. For expression study, Sec13 transcript variant 1, Sec16 transcript variant X12 and CK1α transcript variant 2 were used.

### Sample preparation for MS analysis

Proteins were eluted with 0.1M glycine-HCl (pH3.0), and the eluate was immediately neutralized with 1M Tris-HCl (pH 8.5). Recovered proteins were applied to denaturation, reduction, and alkylation with 8M Urea, 10 mM DTT and 50 mM IAA, respectively. The sample solutions were digested with Lys-C for 3 h and 5-fold diluted with 50 mM ammonium bicarbonate, followed by overnight trypsin digestion. After digestion, the solutions were acidified with 0.5% TFA (final concentration) and desalted with C18-StageTips^31^.

### NanoLC-MS/MS system

NanoLC-MS/MS analyses were conducted by using a timsTOF Pro mass spectrometer equipped with a nanoElute HPLC (Bruker Daltonics, Bremen, Germany). Peptides were separated on an ODYSSEY column (1.6µm C18, 120 Å, 75µm ID, 25 cm) from IonOpticks. The injection volume was 2 µL and the flow rate was 400 nL/min. The mobile phases consisted of (A) 0.1% formic acid and (B) 0.1% formic acid in 80% acetonitrile. A four-step linear gradient of 4–34% B in 90 min, 34–48% B in 15 min, 48–90% B in 5 min, and 90% B for 10 min was employed. The mass scan range was 100–1700 m/z, and the ion mobility was scanned from 0.6 to 1.6 Vs/cm2. The overall scan cycle of 1.16 sec included a single MS scan and ten PASEF MS/MS scans^32^. Low abundance precursor ions below a target intensity value of 20000 counts were repeatedly selected for PASEF-MS/MS. Active exclusion time was set to 0.4 sec.

### Mass spectrometry data analysis

The raw files were analyzed with Fragpipe-MSFragger^33^ for protein identification, and PEAKS studio X+ software (Bioinformatics solutions Inc., Waterloo, ON, CA) with Ascore for phosphosite determination^34^. Peptides and proteins were identified against UniprotKB/Swiss-prot with a precursor mass tolerance of 20 ppm, a fragment ion mass tolerance of 0.05 Da and strict trypsin specificity^35^ allowing for up to 1 missed cleavage. Cysteine carbamidomethylation was set as a fixed modification, and methionine oxidation and protein N-terminal acetylation were allowed as a variable modification. Phosphorylation (STY) were allowed as variable modifications for phosphopeptide identification.

### Immunoprecipitation and western blotting

The experiments were essentially performed as described previously^8^. Cells extracted with extraction buffer consisting of 20 mM Tris-HCl (pH7.4), 100 mM NaCl, 1 mM EDTA, 1% Triton X-100, and protease inhibitors were centrifuged at 20,000 g for 15 min at 4°C. Cell lysates were immunoprecipitated with FLAG M2 antibodies (SIGMA). The beads were washed with TBS/0.1% Triton X-100 for five times followed by elution with DYKDDDK peptide and processed for sample preparation.

### Biotinylated Phos-tag detection

Biotinylated Phos-tag analysis of phosphorylated proteins were conducted essentially as described previously^36^. The samples were resolved by SDS-PAGE followed by blotting to PVDF membranes. The membranes were probed with Phos-tag Biotin (Fuji Film Wako) and Streptavidin-conjugated HRP (Fuji Film Wako) and analyzed using ImageQuant 800 (cytiva).

### siRNA oligos

Stealth select siRNAs for TANGO1S was purchased from Thermo Fisher Scientific. The oligo sequences used were TANGO1S siRNA, 5’-GAAUUGUCGCUUGCGUUCAGCUGUU-3’; For control siRNA, Stealth RNAi™ siRNA negative control med GC Duplex #2 (Thermo Fisher Scientific) was used.

Mission siRNA was purchased from Merck. The oligo sequences used were CK1α siRNA (761), 5’-CAGAAUUUGCGAUGUACUU-3’; CK1α siRNA (131), 5’-CAGUGAAGCUAGAAUCUCA −3’; CK1α siRNA (668), 5’-GGCUAAAGGCUGCAACAAA −3’; CK1δ siRNA (707), 5’-CCAUCGAAGUGUUGUGUAA −3’; CK1ε siRNA (734), 5’-CCUCCGAAUUCUCAACAUA-3’; Sec13 siRNA (3’UTR), 5’-GCACUCAUGUUACGAGGAA-3’; Sec13 siRNA (835), 5’-GUCUCUGGUGGAGACAAUA −3’ For control siRNA, Mission siRNA Universal Negative Control #2 (Merck) was used. The number in the parentheses represents the starting base pair of the target sequence. Silencer select siRNAs for PPP1R15B and PPP6R3 was purchased from Thermo Fisher Scientific. The oligo sequences used were PPP1R15B siRNA(94), 5’-GGCUCUUCUAAGUUCCCGA-3’; PPP1R15B siRNA(2009),5’-GCAGGUUCCAGAAACGAAU-3’; PPP6R3 siRNA (1366), 5’-GGUUACAUGGGACACCUAA-3’; PPP6R3 siRNA (451), 5’-GACCUUAUUAUAAAGCACA-3’;

For control siRNA, Silencer Select Negative Control #1 siRNA (Thermo Fisher Scientific) was used.

### Immunofluorescence microscopy

Immunofluorescence microscopy analysis was performed as described previously^8^. Cells grown on coverslips were washed with PBS, fixed with methanol (6 min at −20°C) or 4% PFA/PBS (10 min at room temperature), and then washed with PBS and blocked in blocking solution (5% BSA in PBS with 0.1% Triton X-100 for 30 min). After blocking, cells were stained with primary antibody (1 h at room temperature) followed by incubation with Alexa Fluor-conjugated secondary antibodies (Alexa Fluor 488, Alexa Fluor 568, and/or Alexa Fluor 647 for 1 h at room temperature). Images were acquired with confocal laser scanning microscopy (Plan Apochromat 63×/1.40 NA oil immersion objective lens; LSM 900 or LSM980; Carl Zeiss). The acquired images were processed with Zen software (Carl Zeiss). All imaging was performed at room temperature.

### RUSH assay

Stable HeLa cells expressing Str-KDEL/ManII-SBP-mCherry under the control of a doxycycline-inducible promoter were seeded onto coverslips. The cells were then treated with 8 ng/ml of doxycycline. After 24 h, biotin was added to the cell culture medium at a final concentration of 40 µM. After 20 min, the cells were fixed with methanol and stained with an anti-GM130 antibody for immunofluorescence. Images were acquired using confocal laser scanning microscopy (LSM 900; Carl Zeiss). Quantification of the percentage of ManII-mCherry signals in the GM130-positive region was then quantified among all ManII-mCherry signals was performed using Fiji-ImageJ.

### FRAP assay

Cells expressing GFP-Sec16 or GFP-Sec16 SA mutant were seeded onto 3.5-cm glass-bottom dishes (Matsunami) and induced with doxycycline 24 hours prior to imaging. Before the assay, the culture medium was replaced with phenol red-free DMEM (Nacalai Tesque). FRAP experiments were performed using confocal laser scanning microscopy (Plan Apochromat 63×/1.40 NA oil immersion objective lens; LSM 980; Carl Zeiss) equipped with an incubation chamber (Tokai Hit, Shizuoka, Japan) with the following parameters: temperature, 37°C; humidity, 100%; and CO_2_ concentration, 5%. Images were acquired and processed with Zen Blue software (RRID:SCR_013672; Carl Zeiss). A circular region of interest (ROI) with a diameter of 2.15 µm was selected and photobleaching was performed using a 488 nm laser at 60% power for 0.51 second. Time-lapse imaging was conducted at a rate of one frame every 0.6304 seconds for a total of 100 frames. Fluorescence intensity within the bleached ROI was quantified using Zen Blue software and normalized to pre-bleach and background intensities. To correct for overall fluorescence decay during image acquisition, fluorescence intensity from a non-bleached ROI was used as a reference for normalization. Recovery curves were fitted and analyzed using Prism 10 (GraphPad).

### In vitro kinase assay

HeLa cells expressing either FLAG-Sec16 1101aa-CT or FLAG-Sec16 1101aa-CT SA mutant were induced with doxycycline for 24 hours, lysed, immunoprecipitated with anti-FLAG antibodies, and eluted with FLAG peptide as described in the section above. The eluates were incubated at 30°C for 1 hour in a reaction buffer added to achieve final concentrations of 20 mM Tris-HCl (pH 7.4), 150 mM NaCl, 0.1% Triton X-100, 5 mM MgClC, and 1 mM [γ-^32^P] ATP. The kinase reaction was stopped by boiling with Laemmli sample buffer. Samples were subjected to SDS-PAGE and subsequently exposed to a phosphor screen, followed by analysis using a Typhoon FLA9500 (GE Healthcare)

### Mitotic Assay

HeLa cells treated with siRNAs were incubated for 22 h and then treated with 2.5 mM thymidine for 16 h. The cells were washed three times with PBS and released for 8 h. Then, the cells were incubated further with 2.5 mM thymidine for 16 h, washed three times with PBS and released for 10 h before fixation. Stained cells were analyzed by confocal laser scanning microscopy (Plan Apochromat 63×/1.40 NA oil immersion objective lens; LSM 900; Carl Zeiss) and processed with Zen software (Carl Zeiss). Intensity scanning and calculating coefficiency were performed by colocalization plugin Coloc2 in Fiji-ImageJ ^37^.

### Statistical analysis

Data were analyzed using an unpaired two-sample Student’s t test for two-group comparisons or a one-way analysis of variance followed by Tukey’s test or Dunnett’s test for multiple comparisons using GraphPad Prism software (RRID:SCR_002798; GraphPad). Data distribution was assumed to be normal, but this was not formally tested. All graphs were created using GraphPad Prism software (RRID:SCR_002798; GraphPad).

## ACKNOWLEDGEMENTS

This work was supported by JSPS Grants-in-Aid for Scientific Research (22H02760 and 23K24023 to MM, 24K18067 to MA, 23H05254 to YK and 23H02430, 23K27123, and 24K22065 to KS) from the Ministry of Education, Culture, Sports, Science, and Technology of Japan and Naito Foundation (KS and MM). KS received support from the Takeda Science Foundation, Asahi Glass Foundation, and Princess Takamatsu Cancer Research Foundation.

## AUTHOR CONTRIBUTIONS

M.M. designed and performed research, analyzed data, and wrote the manuscript; M.A. performed research, analyzed data; M.W. performed research, analyzed data; Y.K. performed research; and K.S. designed and performed research, analyzed data, and wrote the manuscript.

---

**Fig. S1. ERES-localized Sec16 signal was markedly reduced upon Sec13 knockdown.** HeLa cells were transfected with indicated siRNAs. Cells were then fixed with cold methanol and stained with anti-Sec16-C or anti-TANGO1-CT antibodies. Bars, 10 µm.

**Fig. S2. CK1**α **Phosphorylates Sec16.** 293T cells were transfected with FLAG-tagged Sec16 (1101–1890 aa) with HA-tagged kinases as indicated. Cell lysates were immunoprecipitated with anti-FLAG antibody and eluted with a FLAG peptide. Eluates were then subjected to SDS-PAGE followed by biotinylated Phos-tag detection or western blotting with anti-FLAG antibody. Lysates were subjected to SDS-PAGE followed by western blotting with anti-HA antibody.

**Fig. S3. Sec16 localizes to ERES regardless of its phosphorylation status.** Doxycycline-inducible HeLa cells expressing either GFP-Sec16, GFP-Sec16 SA, GFP-Sec16 SE were fixed and stained with TANGO1-CT antibody. Bars, 10 µm.

**Fig. S4. CK1α inhibitor D4476 reduces the turnover rate of Sec16 at ER exit site.** (A) Doxycycline-inducible HeLa cells expressing GFP-Sec16 were treated with or without 100 µM D4476 for 4 h and subjected to FRAP assay. Representative images are shown before bleaching (pre-bleaching) and at the indicated time points after photobleaching. Bars, 1 µm. (B) Quantification of fluorescence recovery from FRAP assays as in (A). FRAP curves show the mean fluorescence intensity of 10 individual regions and error bars indicate the S.D. Fluorescence values were normalized to the pre-bleach intensity.

**Fig. S5. FAM83D, FAM83G and FAM83H do not colocalize with Sec16 when co-expressed with CK1**α **kinase-dead mutant.** HeLa cells transfected with FLAG-FAM83D or FLAG-FAM83G or FLAG-FAM83H with HA-CK1α K46A were fixed and stained with anti-Sec16-C, anti-FLAG, and anti-HA antibodies. Bars, 10 µm.

**Fig. S6. RUSH assay shows that wild-type TANGO1S, but not the 21SA mutant, rescues secretion defects upon TANGO1S knockdown.** Doxycycline-inducible HeLa cells expressing either TANGO1S-FLAG (A) or TANGO1S SA mutant-FLAG (B) were transfected with the indicated siRNAs. The cells were treated with or without doxycycline and transfected with Str-KDEL_ManII-SBP-mCherry. RUSH chase was initiated with the addition of biotin and terminated after 20 min with cold methanol fixation. The cells were stained with anti-GM130 and anti-FLAG antibodies. Bars, 10 μm.

**Fig. S7. pSec16 antibody detects decreased Sec16 phosphorylation upon Sec13 knockdown.** Doxycycline-inducible HeLa cells expressing FLAG-tagged Sec16 (1101aa–CT) were transfected with indicated siRNAs. Cell lysates were immunoprecipitated with anti-FLAG antibody and eluted with a FLAG peptide. Eluates were subjected to SDS-PAGE followed by western blotting with anti-FLAG and anti-pSec16 antibodies. Lysates were subjected to SDS-PAGE followed by western blotting with anti-Sec13 and anti-GAPDH antibodies.

**Fig. S8. CK1**α **is also required for mitotic ER exit site disassembly.** HeLa cells transfected with the indicated siRNAs were synchronized at mitosis using a double thymidine block, then fixed and stained with anti-Sec16 and anti-Sec31 antibodies. Pearson’s colocalization coefficient between Sec16 and Sec31 was quantified. Each circle represents an individual cell (n = 15); red bars indicate the mean and 95% confidence interval. Data were analyzed by one-way ANOVA and Dunnett’s multiple comparison test.

