## Supplemental for "Phosphorylation-Coupled Autoregulation Maintains Functional ER Exit Sites"

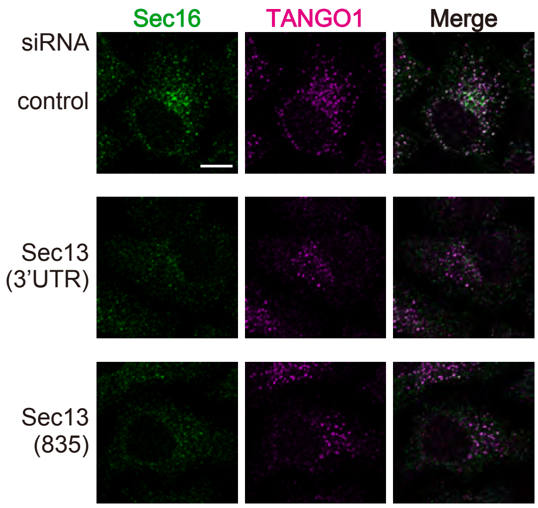


Supplemental Figure 1

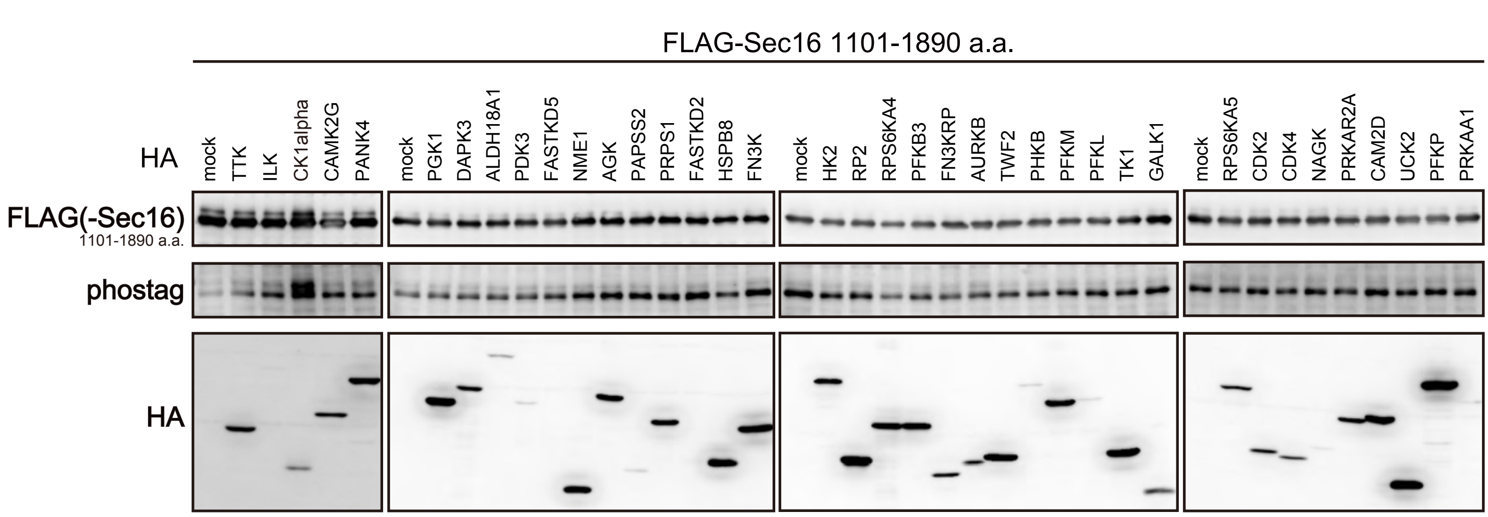


Supplemental Figure 2

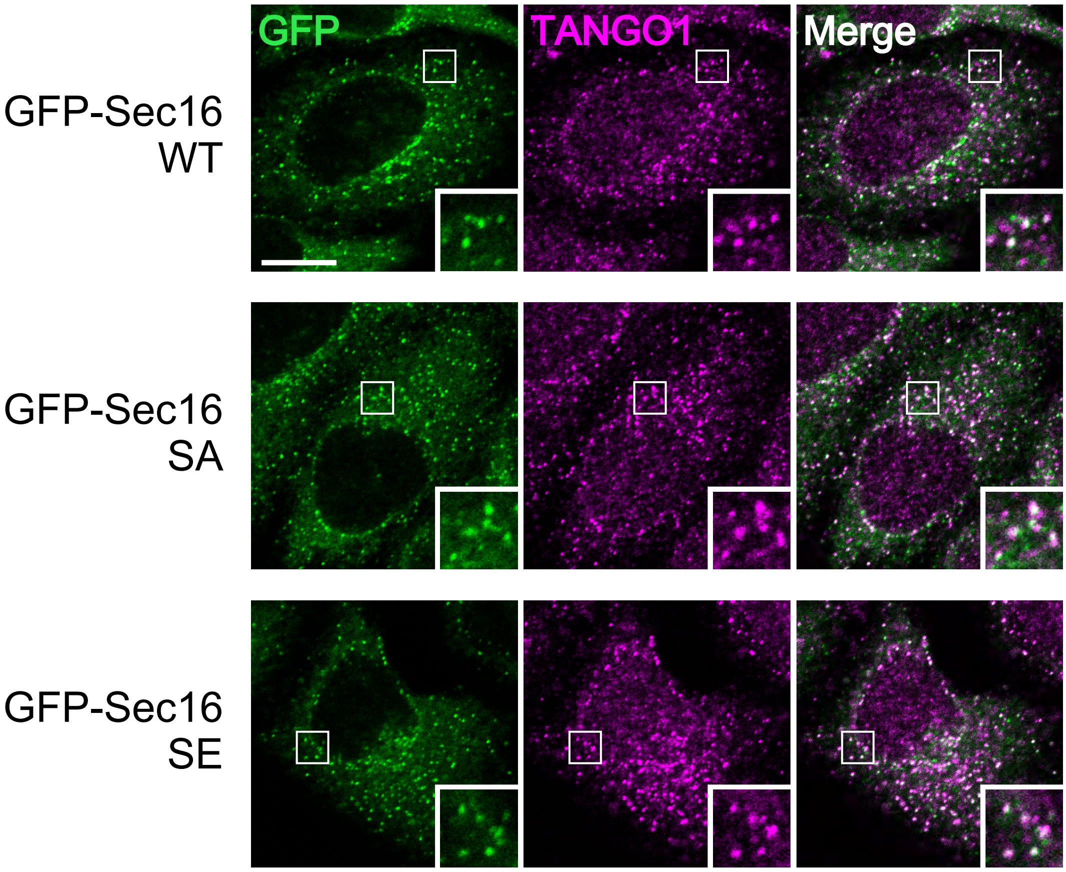


Supplemental Figure 3

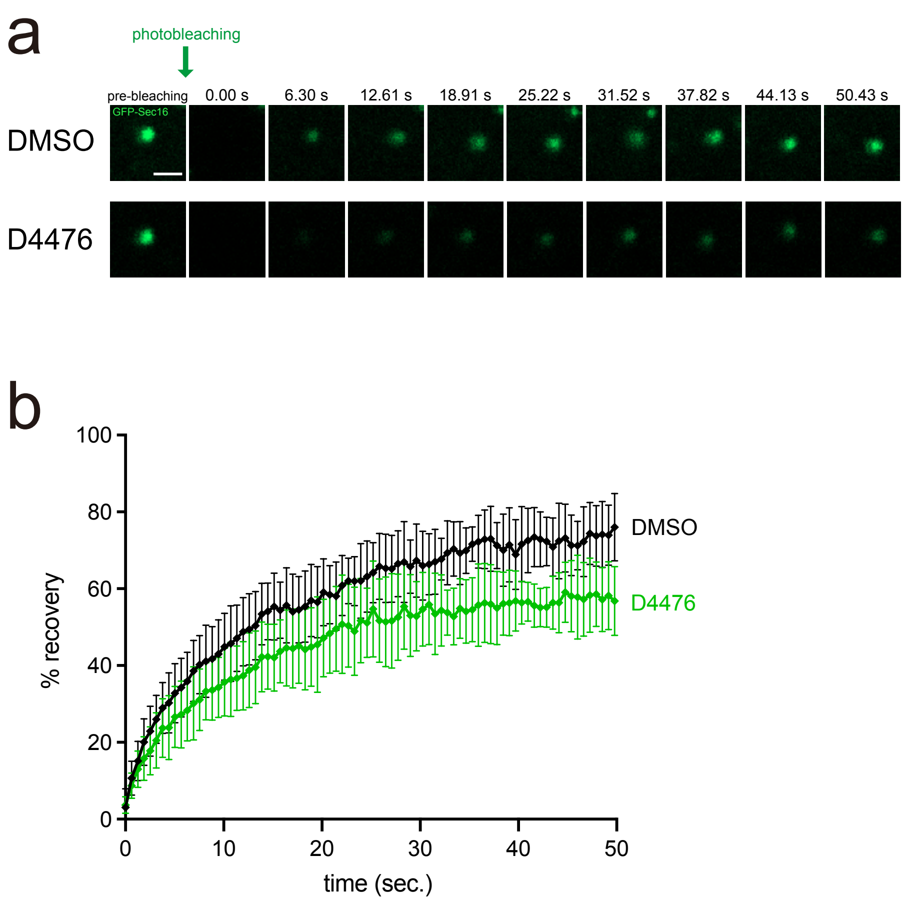


Supplemental Figure 4

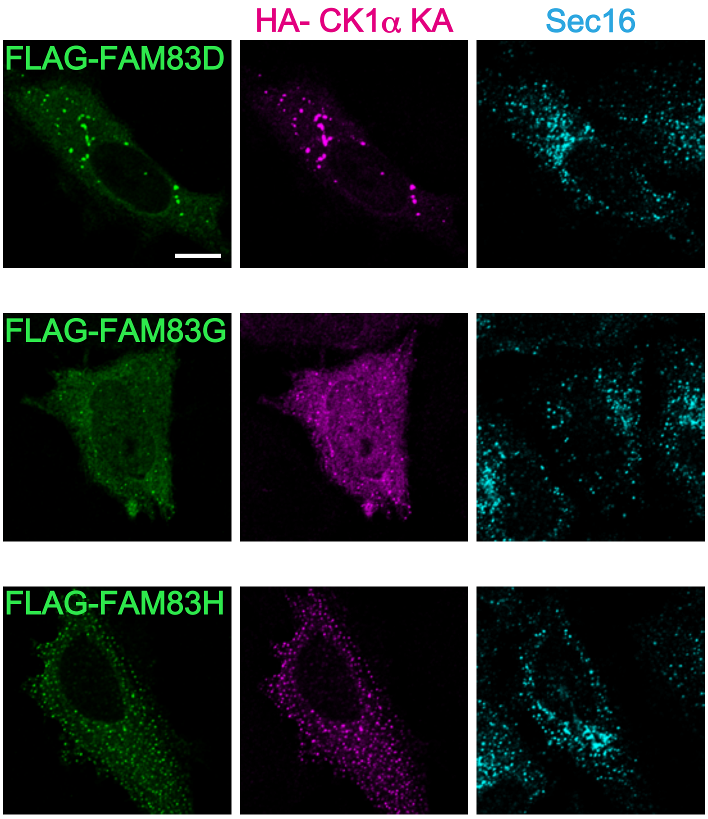


Supplemental Figure 5

a

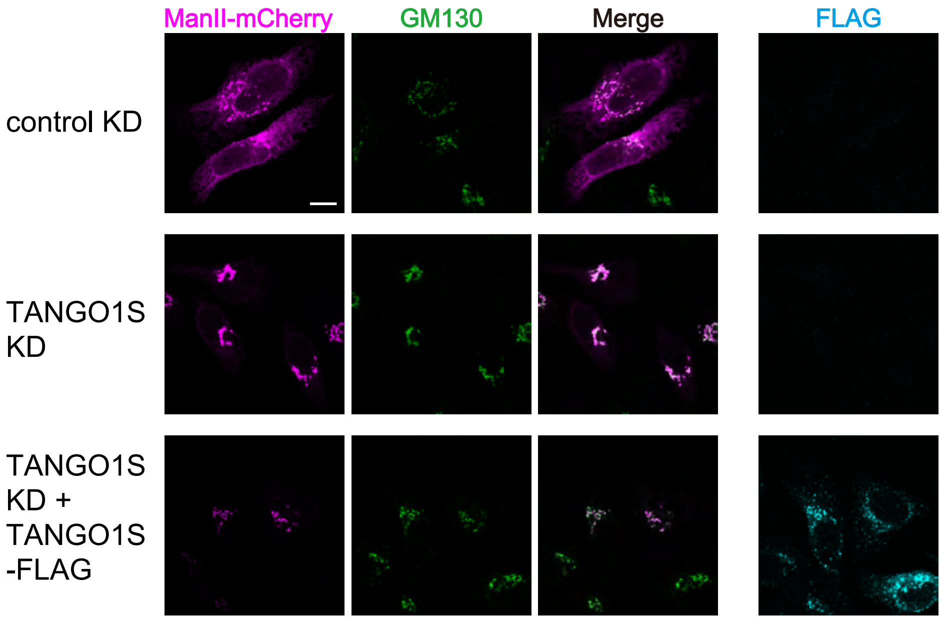


b

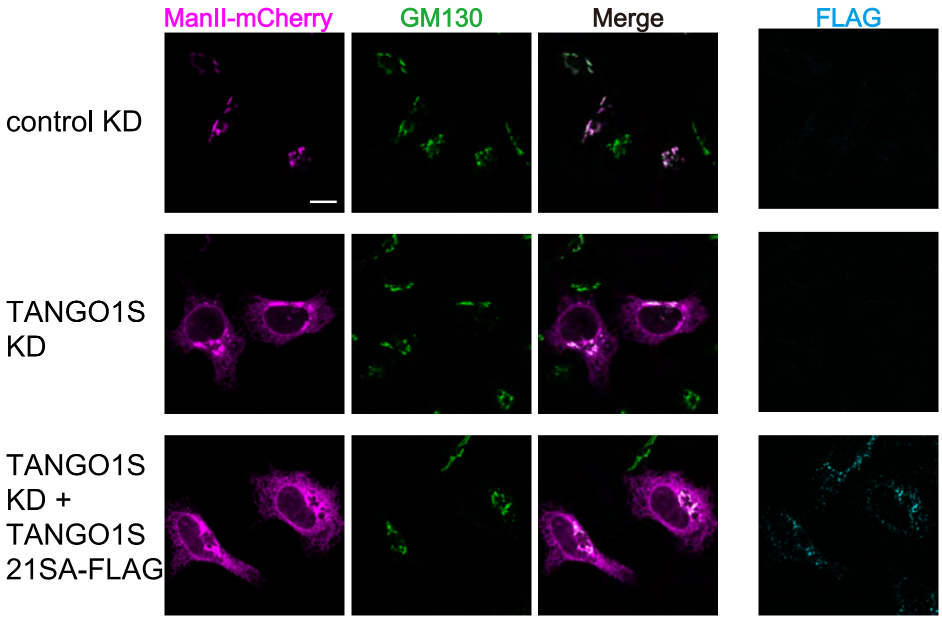


Supplemental Figure 6

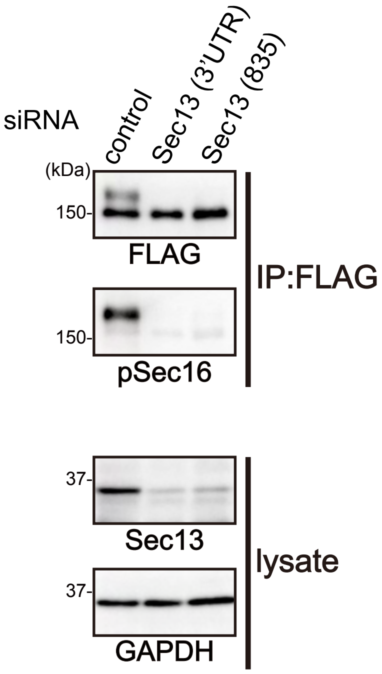


Supplemental Figure 7

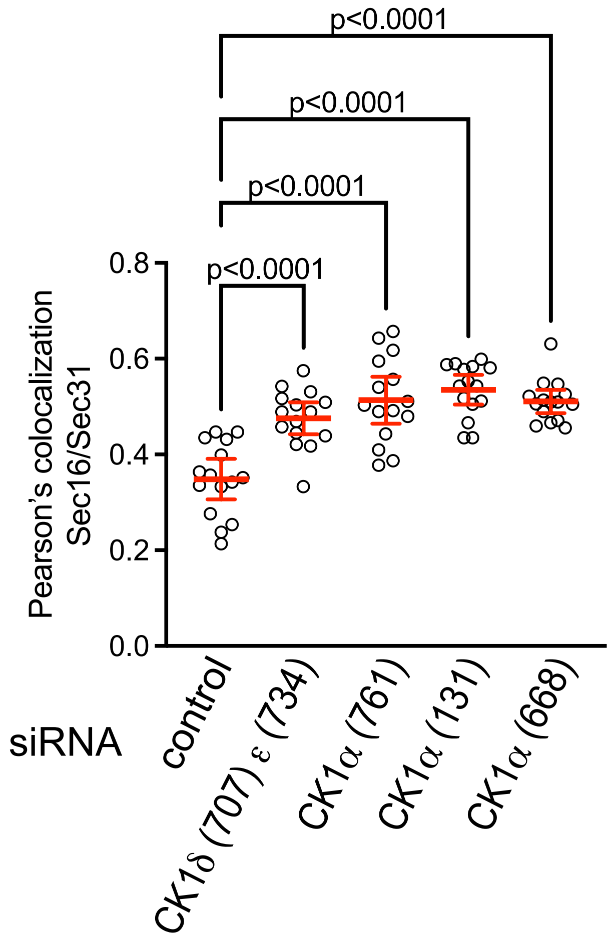


Supplemental Figure 8
